# Genotype-Informed *Drosophila* Kinome Screening Prioritizes *PRKX* as a Candidate Vulnerability in Bladder Cancer

**DOI:** 10.64898/2026.11.08.765218

**Authors:** Takuya Moriguchi, Chihiro Kamijo, Taku Kimura, Taiga Hirata, Reo Sato, Ryodai Yamamura, Takahiro Osawa, Nobuo Shinohara, Takashige Abe, Masahiro Sonoshita

## Abstract

Muscle-invasive bladder cancer is molecularly heterogeneous, and the basal/squamous subtype frequently exhibits elevated *EGFR* expression and *TP53* and *RB1* alterations. Functional vulnerabilities associated with these alterations remain incompletely defined, while large-scale *in vivo* screening in mammalian models is experimentally demanding. Therefore, we developed a genotype-defined *Drosophila melanogaster* model combining *Egfr* overexpression, mutant *p53*, and *Rbf* suppression in Malpighian tubules. The resulting ‘*3-hit’* model showed increased Malpighian tubule volume and reduced adult eclosion rate. A dominant modifier screen targeting 216 kinase genes identified 73 genes whose perturbation significantly increased the eclosion rate, and a complementary RNA interference screen identified eight kinases whose knockdown increased the eclosion rate. Among these hits, *Pka-C3*, a *Drosophila* ortholog of human *PRKX*, emerged as one of the strongest suppressors of the reduced adult eclosion rate in the *3-hit* model. *PRKX* was prioritized for further study because public transcriptomic datasets showed higher *PRKX* expression in bladder tumors than in normal bladder tissue. *PRKX* knockdown reduced spheroid volume in four human bladder cancer cell lines with distinct *EGFR*, *TP53*, and *RB1* profiles. Both small interfering RNAs produced significant reductions in two cell lines, whereas only one siRNA reached significance in the other two. These findings nominate *PRKX* as a candidate vulnerability in bladder cancer and highlight a cross-species approach that combines genotype-informed *Drosophila* screening with functional testing in human cancer cells to prioritize targets for further investigation.

## Introduction

Bladder cancer accounted for approximately 614,000 new cases worldwide, ranking ninth among all cancers ^1^. Clinically, bladder cancer is broadly classified as non-muscle-invasive bladder cancer (NMIBC) or muscle-invasive bladder cancer (MIBC) according to invasion of the muscularis propria. Whereas NMIBC is generally managed with bladder-preserving treatment, MIBC often requires multimodal treatment combining local and systemic therapies ^2–4^. Outcomes and treatment responses vary substantially among patients carrying MIBC with similar clinicopathological features, reflecting the biological heterogeneity of the disease. Genomic and transcriptomic studies have identified molecular subtypes with distinct differentiation programs, oncogenic alterations, and tumor microenvironments, providing a framework for investigating therapeutic targets within defined molecular contexts ^5–7^.

The consensus molecular classification defines six MIBC classes: luminal papillary, luminal non-specified, luminal unstable, stroma-rich, basal/squamous, and neuroendocrine-like. The basal/squamous subtype of bladder cancer (BSBC) was the largest class, accounting for approximately 35% of MIBC in the consensus dataset and was associated with poorer overall survival compared with other classes ^8^. BSBC is characterized by basal and squamous differentiation programs and relatively low expression of luminal markers ^8, 9^, together with *EGFR* overexpression ^10^. Alterations in the *TP53* and *RB1* tumor-suppressor genes are also frequent; analysis of The Cancer Genome Atlas (TCGA) exome data within the consensus study identified *TP53* and *RB1* mutations in approximately 61% and 25% of BSBC, respectively ^7, 8, 10^. These features provide a rationale for investigating therapeutic vulnerabilities in genetic contexts associated with BSBC.

Molecular profiling identifies recurrent alterations, but functional perturbation is needed to determine which genes modify phenotypes arising from these alterations. *Drosophila melanogaster* provides a genetically tractable whole-animal platform in which multiple oncogenic alterations can be introduced into a defined tissue and combined with dominant modifier or RNAi screening ^11–15^. Such models have enabled the identification of signaling dependencies and pharmacological vulnerabilities for subsequent evaluation in mammalian cancer systems ^16–21^. Using this approach, we previously developed and used *Drosophila* models of genetic alterations observed in pancreatic ductal adenocarcinoma (PDAC) to identify kinase and metabolic vulnerabilities and evaluate prioritized candidates in human cancer models ^22–25^. This strategy provides a basis for testing modifiers of selected BSBC-associated alterations. Kinases are particularly attractive candidates because they regulate cancer-associated signaling and include established targets for anticancer drugs ^26^.

Here, we developed a genotype-defined *Drosophila* model combining *Egfr* overexpression, mutant *p53*, and *Rbf* suppression to represent selected ‘*3-hit*’ BSBC-associated molecular alterations. Using adult eclosion rate as a whole-animal phenotypic readout, we screened for kinase genes whose perturbation rescued the reduced eclosion phenotype of this model. Among the strongest suppressors was *Pka-C3*, a *Drosophila* ortholog of *human protein kinase X* (*PRKX*), whose heterozygosity significantly rescued the reduced adult eclosion rate. Integration with human bladder tumor-expression data prioritized *PRKX* for functional evaluation. *PRKX* knockdown reduced spheroid volume in multiple human bladder cancer cell lines, supporting its further investigation as a candidate vulnerability in bladder cancer.

## Materials and Methods

### Construction of flies modeling key genetic features of BSBC

Genomic DNA was extracted using the DNeasy Blood & Tissue Kit (QIAGEN, Hilden, Germany) from two transgenic *Drosophila* lines obtained from the Bloomington *Drosophila* Stock Center (BDSC; Bloomington, IN, USA): *UAS*-*Egfr* (BDSC #5368) and *UAS*-*p53*^R259H^ (BDSC #6582). The *p53*^R259H^ substitution corresponds to the human *TP53*^R273H^ hotspot mutation^27^. The full-length *Egfr* and *p53*^R259H^ cDNA sequences were amplified by PCR from the respective genomic DNA templates. The PCR products were cloned into the pCR-Blunt II-TOPO vector using the Zero Blunt TOPO PCR Cloning Kit (Invitrogen, Carlsbad, CA, USA). We performed site-directed mutagenesis using PfuTurbo DNA polymerase (Agilent Technologies, Santa Clara, CA, USA), followed by DpnI digestion, based on the QuikChange site-directed mutagenesis protocol. We first reverted the *p53*^R259H^ substitution to the wild-type sequence, then introduced the *p53*^R234Q^ substitution, corresponding to the human *TP53*^R248Q^ hotspot mutation ^27^. Each mutagenesis step was verified by sequencing. The resulting *p53*^R234Q^ construct contained a C-terminal c-Myc tag.

The *Egfr* and *p53*^R234Q^ fragments were subcloned into a multigenic expression vector containing two upstream activation sequence (*UAS*) cloning cassettes and one short hairpin RNA (shRNA) cloning cassette ^28, 29^. The resulting plasmid was injected into *y¹ w⁶⁷ᶜ²³; P{CaryP}attP40* embryos to generate *UAS-Egfr,UAS-p53*^R234Q^ flies. An shRNA targeting *Rbf*, the *Drosophila* ortholog of human *RB1*, with the target sequence CGAGGTGTTCATCGACATCAA was cloned into the expression vector, and the resulting plasmid was injected into *y¹ w⁶⁷ᶜ²³; P{CaryP}attP16* embryos to generate *UAS-Rbf*^shRNA^ flies. Both transgenic lines carried insertions on the second chromosome and were generated by BestGene Inc. (Chino Hills, CA, USA). The lines were crossed, and meiotic recombination was used to combine the transgenes and generate *UAS-Egfr,UAS-p53*^R234Q^*,UAS-Rbf*^shRNA^ (hereafter referred to as *UAS*-*3-hit*) flies.

The *c507-GAL4* (*w[*]; P{w[+mW.hs]=GawB}Alp4*[*c507*], BDSC #30840) was used to express the transgenes in Malpighian tubules (MTs). The *c507-GAL4* and *UAS*-*GFP* (*y[*] w[*]; P{w[+mC]=UAS-2xEGFP}AH3*, BDSC #6658) transgenes were combined by meiotic recombination to generate the *c507-GAL4*,*UAS*-*GFP* line (hereafter referred to as *c507*>*GFP*). This line was crossed with *UAS-3-hit* flies to generate *UAS-3-hit*;*c507-GAL4*,*UAS-GFP* flies (hereafter referred to as *c507>3-hit*), which coexpressed GFP and the three modeled genetic alterations in MTs. For MT phenotyping, *c507*>*GFP* larvae served as controls.

### Malpighian tubule analysis

MTs were dissected from first-instar (L1) larvae maintained at 29°C, fixed in 4% paraformaldehyde (PFA; Nacalai Tesque, Kyoto, Japan) for 20 min, and washed three times for 10 min each in phosphate-buffered saline (PBS; Fujifilm Wako, Osaka, Japan). Samples were mounted in Vectashield Vibrance Antifade Mounting Medium containing 0.9 µg/mL 4′,6-diamidino-2-phenylindole (DAPI; Vector Laboratories, Malvern, PA, USA). Images were acquired using a Leica DMi8 fluorescence microscope (Leica, Wetzlar, Germany) at 2-µm intervals along the z-axis. GFP-positive areas were measured using ImageJ (US National Institutes of Health, Bethesda, MD, USA).

GFP-positive MT volume was estimated by multiplying the GFP-positive area in each image (µm²) by the z-interval (2 µm) and summing across the image series. Five larvae, corresponding to five MT pairs, were analyzed per group, with each larva treated as a biological replicate (*n* = 5).

### Genomic PCR validation of transgenes

Genomic DNA was extracted from *UAS*-*3-hit* larvae using the DNeasy Blood & Tissue Kit. Genomic PCR was performed to confirm the presence of the *Egfr* and *p53*^R234Q^ transgenes. For *Egfr*, the forward primer was designed to span the vector–*Egfr* insert junction, allowing specific amplification of the transgene rather than the endogenous *Egfr* locus. For *p53*^R234Q^, the reverse primer contained sequence corresponding to the C-terminal c-Myc tag, enabling transgene-specific amplification. The expected PCR product sizes were 561 bp for *Egfr* and 368 bp for *p53*^R234Q^. The corresponding plasmids containing the *Egfr* or *p53*^R234Q^ transgene were used as positive controls. PCR products were separated by electrophoresis on a 1% agarose gel in Tris-acetate-EDTA buffer. Gene Ladder Wide 1 (Nippon Gene, Tokyo, Japan) was used as a DNA size marker. Primer sequences are provided in Supplementary Table S1.

### *Drosophila* stocks and genetic screening

Kinase-mutant and *UAS*-RNAi stocks were obtained from the BDSC, KYOTO Stock Center (Kyoto, Japan), and Vienna *Drosophila* Resource Center (VDRC; Vienna, Austria). Alleles, RNAi transgenes, and stock identifiers are listed in Supplementary Table S2. Stocks carrying *FM6*, *FM7a*, *FM7c*, or *FM7i* balancers were rebalanced with *FM7c-Tb-RFP* (BDSC #36337); those carrying *CyO*, *SM5*, or *SM6a* were rebalanced with *CyO-Tb-RFP* (BDSC #36336); and those carrying *TM3*, *TM6C*, or *MKRS* were rebalanced with *TM6B* carrying *Tubby* (*Tb*) as a visible marker. A *GAL80*-bearing balancer was used to suppress *GAL4*-dependent expression of the *UAS*-*3-hit* transgenes during stock maintenance ^30^.

Genetic screening was performed as previously described ^22^, with modifications for the *c507*>*3-hit* flies. The dominant modifier screen covered 216 genes. Crossing schemes are shown in Supplementary Fig. S1. For the dominant modifier screen of X-linked genes, *3-hit* males were crossed with *white^−^* control females or kinase-mutant females. For genes on chromosomes 2, 3, or 4, *3-hit* females were crossed with *white−* control males or kinase-mutant males. For the complementary RNAi screen, *3-hit* females were crossed with males carrying *UAS*-kinase RNAi transgenes.

Progeny were reared on standard fly food at 29°C for 10 days. Only non-*Tb* progeny were included in the pupal and adult counts. Adult eclosion rate was calculated for each cross as 100 × (number of eclosed non-*Tb* adults/number of non-*Tb* pupae). For the X-linked screen, female and male progeny were pooled and analyzed together. Each screening condition and its corresponding control were assessed in three independent crosses. Changes in adult eclosion rate were expressed in percentage points relative to the respective control. Human orthologs were predicted using the DRSC Integrative Ortholog Prediction Tool (DIOPT) ^31^. Fly food preparation is described in the Supplementary Information.

### Public datasets and cell-line annotation

Expression of *TAF1*, *TTBK2*, and *PRKX* in human bladder tumors and normal bladder tissues was compared using Gene Expression Profiling Interactive Analysis 2 (GEPIA2) ^32^. Analyses used TCGA and GTEx data and included 404 tumor samples and 28 normal tissue samples. Expression was displayed as log_2_(TPM+1), where TPM denotes transcripts per million. Tumor and normal tissue groups were compared by one-way analysis of variance (ANOVA) as implemented in GEPIA2. Cell line annotations were compiled from Cellosaurus ^33^ and cross-referenced with the Cancer Dependency Map (DepMap) ^34^ and Cancer Cell Line Encyclopedia (CCLE) ^35^. *EGFR* expression and *TP53* and *RB1* alterations are summarized in Supplementary Table S3.

### Cell culture

Human bladder cancer cell lines 5637, UM-UC-3, HT-1376, and HT-1197 were obtained from the American Type Culture Collection (ATCC, Manassas, VA, USA). Cells were maintained at 37°C in a humidified atmosphere containing 5% CO_2_ in the appropriate media supplemented with 10% fetal bovine serum (Thermo Fisher Scientific) and 1% penicillin/streptomycin (Nacalai Tesque, Kyoto, Japan). Culture media and cell line RRIDs are provided in Supplementary Table S4. Cells were used within 2 months after thawing.

### siRNA transfection

Cells were seeded in 6-well plates and transfected 24 h after seeding with 5 nM si*PRKX*-targeting siRNA or non-silencing control siRNA using Lipofectamine RNAiMAX (Thermo Fisher Scientific), according to the manufacturer’s instructions. Two siRNAs targeting *PRKX*, s11195 (hereafter designated as si*PRKX* #1) and s224411 (si*PRKX* #2), and Silencer Select Negative Control No. 1 siRNA (siNs) were purchased from Thermo Fisher Scientific. Knockdown efficiency was assessed by RT-qPCR 48 h after transfection.

### Spheroid formation and volume measurement

At 48 h after transfection, cells were trypsinized, counted, and reseeded at 1,500 cells in 200 µL per well into U-bottom, cell-repellent 96-well plates (Greiner Bio-One, Kremsmünster, Austria). Spheroids were imaged on day 7 after reseeding, and their major and minor axes were measured. Volume was estimated as *V* = (π/6) × *L* × *W*², where *L* and *W* denote the major and minor axes, respectively. Each condition included 3–4 technical replicate wells, with one spheroid analyzed per well. Within each independent experiment, spheroid volumes from the replicate wells were averaged for each condition, yielding one mean value per condition per experiment. These experiment-level means from three independent experiments (*n* = 3) were used for statistical analysis (Fig. 3C). The three siRNA conditions within each independent experiment were treated as matched. Statistical significance was assessed using repeated-measures one-way ANOVA followed by Dunnett’s multiple-comparisons test, with siNs as the control group.

### RNA extraction and RT-qPCR

For *Drosophila* samples, 10–20 MTs from L1 larvae of each indicated genotype were pooled, and total RNA was extracted using the RNeasy Mini Kit (QIAGEN). For human bladder cancer cells, total RNA was extracted 48 h after siRNA transfection using Isogen II (Nippon Gene, Tokyo, Japan). RNA (20–100 ng) was reverse-transcribed using ReverTra Ace qPCR RT Master Mix (Toyobo, Osaka, Japan) at 37°C for 15 min, followed by 50°C for 5 min and 98°C for 5 min.

Quantitative PCR was performed using THUNDERBIRD SYBR qPCR Mix (Toyobo) on a StepOnePlus Real-Time PCR System (Thermo Fisher Scientific). Cycling conditions were 95°C for 1 min, followed by 40 cycles of 95°C for 15 s and 60°C for 45 s. *RpL32* (*ribosomal protein L32*) and *ACTB* (*β-actin*) served as reference genes for *Drosophila* and human samples, respectively. Relative expression was calculated using the 2^-ΔΔCt^ method ^36^ and expressed relative to the corresponding control. Primer sequences are listed in Supplementary Table S1.

### Statistical analysis

Statistical analyses were performed using GraphPad Prism version 11.1.0 (GraphPad Software, Boston, MA, USA) and EZR version 1.68 (Saitama Medical Center, Jichi Medical University, Saitama, Japan), a graphical user interface for R version 4.5.1 ^37^. Data are presented as mean ± standard deviation (SD), unless otherwise indicated. Sample sizes and experimental units are specified in the corresponding assay descriptions and figure legends.

GFP-positive MT volume was compared using an unpaired, two-tailed Welch’s *t*-test. *Rbf* expression in MTs was compared using an unpaired, two-tailed Student’s *t*-test. Temperature-dependent differences in adult eclosion rate were analyzed by ordinary two-way ANOVA followed by Sidak’s multiple-comparisons test, comparing *c507>GFP* and *c507>3-hit* flies at each temperature.

For the dominant modifier screen, X-linked and autosomal screens were analyzed separately because they used different crossing schemes and corresponding controls. Each screen was analyzed by one-way ANOVA followed by Dunnett’s multiple-comparisons test against its respective control. Genes whose perturbation increased adult eclosion rate with a Dunnett’s-adjusted *P* value < 0.05 were classified as significant suppressors. The complementary RNAi screen was analyzed by one-way ANOVA followed by Dunnett’s multiple-comparisons test against the control cross. *PRKX* RT-qPCR data were analyzed by ordinary one-way ANOVA followed by Dunnett’s multiple-comparisons test, with siNs as the control group. Spheroid-volume data were analyzed by repeated-measures one-way ANOVA followed by Dunnett’s multiple-comparisons test, with siNs as the control group.

## Results

### Establishment and phenotypic characterization of a *Drosophila* model of key BSBC-associated genetic alterations

To model selected BSBC-associated molecular alterations, we combined *Egfr* overexpression, mutant *p53*^R234Q^, and *Rbf* suppression in *Drosophila*, generating a ‘*3-hit*’ model (Fig. 1A). Genomic PCR confirmed the presence of the *Egfr* and *p53*^R234Q^ transgenes, and RT-qPCR confirmed reduced endogenous *Rbf* expression (Supplementary Fig. S2). We expressed these transgenes in MTs, a genetically tractable renal epithelium with conserved transport functions ^38, 39^ (Fig. 1B). The *c507-GAL4* driver induced the *UAS-3-hit* transgenes together with *UAS-GFP*, which marked the transgene-expressing domain ^11, 12^. Compared with *c507>GFP* controls, *c507>3-hit* L1 larvae exhibited an expanded GFP-positive MT domain (Fig. 1C). Quantification showed a significant increase in GFP-positive MT volume in the *c507>3-hit* group (Fig. 1D).

**Figure 1.**
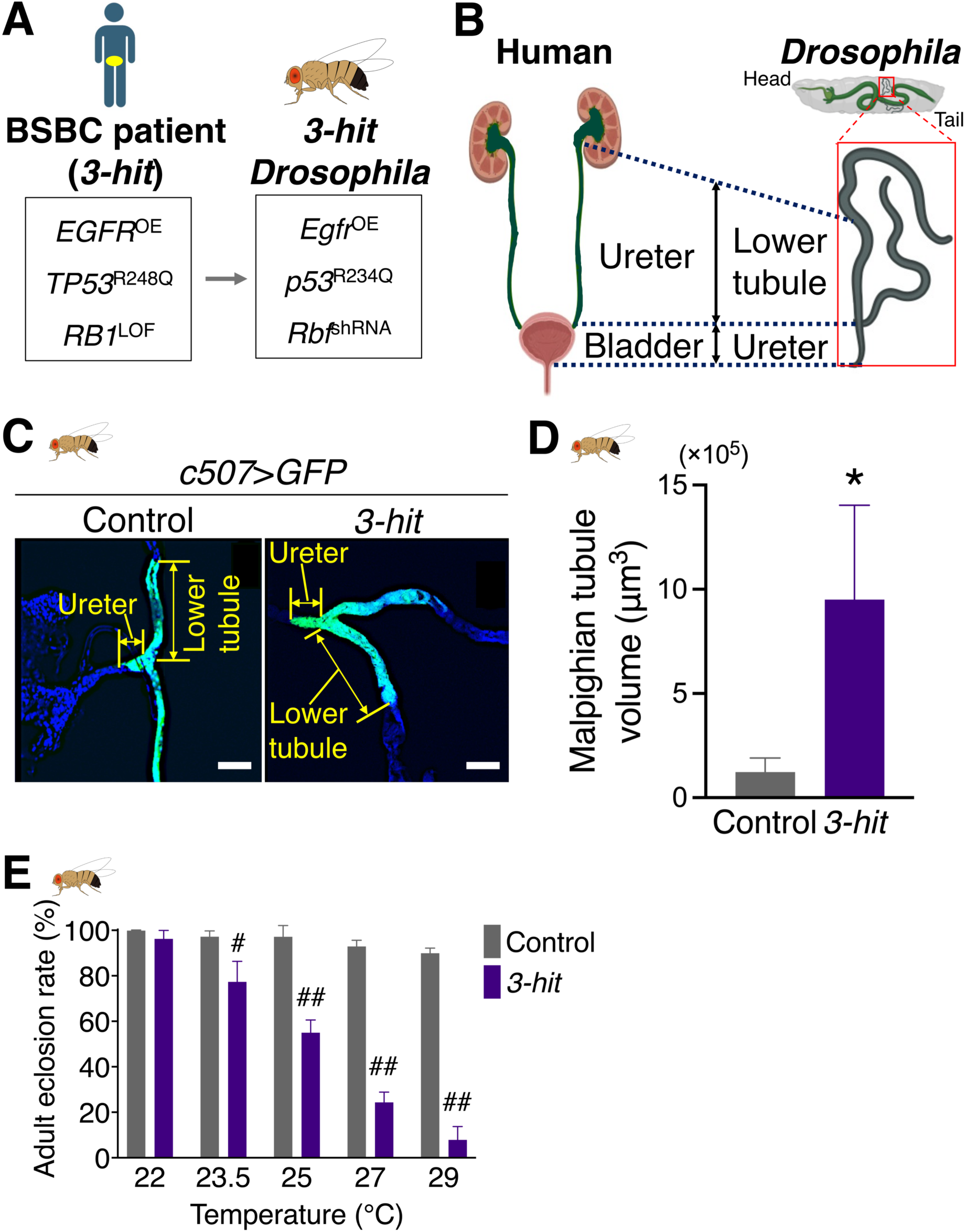
A *Drosophila* model of selected basal/squamous bladder cancer-associated alterations exhibits increased Malpighian tubule volume and reduced adult eclosion rate. **(A)** Schematic of the human genetic alterations represented in the *3-hit Drosophila* model. *EGFR* overexpression, the *TP53*^R248Q^ mutation, and *RB1* loss are modeled by *Egfr* overexpression (OE), *p53*^R234Q^ mutation, and *Rbf* knockdown, respectively. BSBC, basal/squamous subtype of bladder cancer; LOF, loss of function. **(B)** Schematic of the human urinary system and the *Drosophila* excretory system, highlighting the lower tubule and ureter region of the Malpighian tubules (MTs) examined in this study alongside the human ureter and bladder for anatomical comparison. MTs provide a genetically tractable renal/excretory epithelium for modeling the selected molecular alterations. **(C)** Representative fluorescence images of MTs from control (*c507*>*GFP*) and *3-hit* (*c507*>*3-hit*) L1 larvae. In *3-hit* larvae, *c507*-*GAL4* drives *GFP* together with *Egfr*, *p53*^R234Q^, and *Rbf*^shRNA^. GFP (green) marks the transgene-expressing domain, and DAPI (blue) labels nuclei. Scale bars, 100 µm. **(D)** GFP-positive MT volume (μm^3^). Five larvae, corresponding to five MT pairs, were analyzed per group (*n* = 5 biological replicates). Bars show mean ± SD. \**P* < 0.05 by an unpaired, two-tailed Welch’s *t*-test. Transgene validation and *Rbf* expression analysis are shown in Supplementary Figure S2. **(E)** Adult eclosion rate of control and *3-hit* flies reared at 22, 23.5, 25, 27, or 29°C. Eclosion rate was calculated as 100 × (number of eclosed non-*Tb* adults/number of non-*Tb* pupae). Bars show mean ± SD from three biological replicates per condition. ^#^*P* < 0.05, ^##^*P* < 0.0001 versus the corresponding control at the same temperature by ordinary two-way ANOVA followed by Sidak’s multiple-comparisons test.

To establish a sensitized reduced-eclosion phenotype suitable for genetic modifier screening, we examined adult eclosion rates across rearing temperatures from 22 to 29°C, taking advantage of the temperature dependence of *GAL4*-mediated transgene expression. The adult eclosion rate of control flies remained high across this range, whereas that of *c507*>*3-hit* flies progressively decreased with increasing temperature, reaching approximately 50% at 25°C and less than 10% at 29°C (Fig. 1E). The adult eclosion rate was significantly lower in *c507*>*3-hit* flies than in the corresponding controls at 23.5, 25, 27, and 29°C. The low but measurable adult eclosion rate at 29°C provided a sensitized background in which genetic suppression of the *3-hit* phenotype could be detected. Therefore, 29°C was used for subsequent genetic screening. Together, these findings established increased GFP-positive MT volume and a reduced adult eclosion rate as tissue-level and whole-animal phenotypes of the *3-hit* model.

### Whole-animal kinase screening and tumor expression analysis prioritize *PRKX*

To identify genetic modifiers of the *3-hit* phenotype, we performed a kinase modifier screen using mutant lines targeting 216 *Drosophila* genes (Fig. 2A; Fig. S1 and Table S2). Adult eclosion rate was defined as the percentage of non-*Tb* pupae that eclosed as adults. Control crosses between *3-hit* flies and *white*^−^ flies yielded an adult eclosion rate of approximately 75% under the screening conditions. Because the X-linked and autosomal screens used different crossing schemes, each screen was analyzed separately against its respective control.

**Figure 2.**
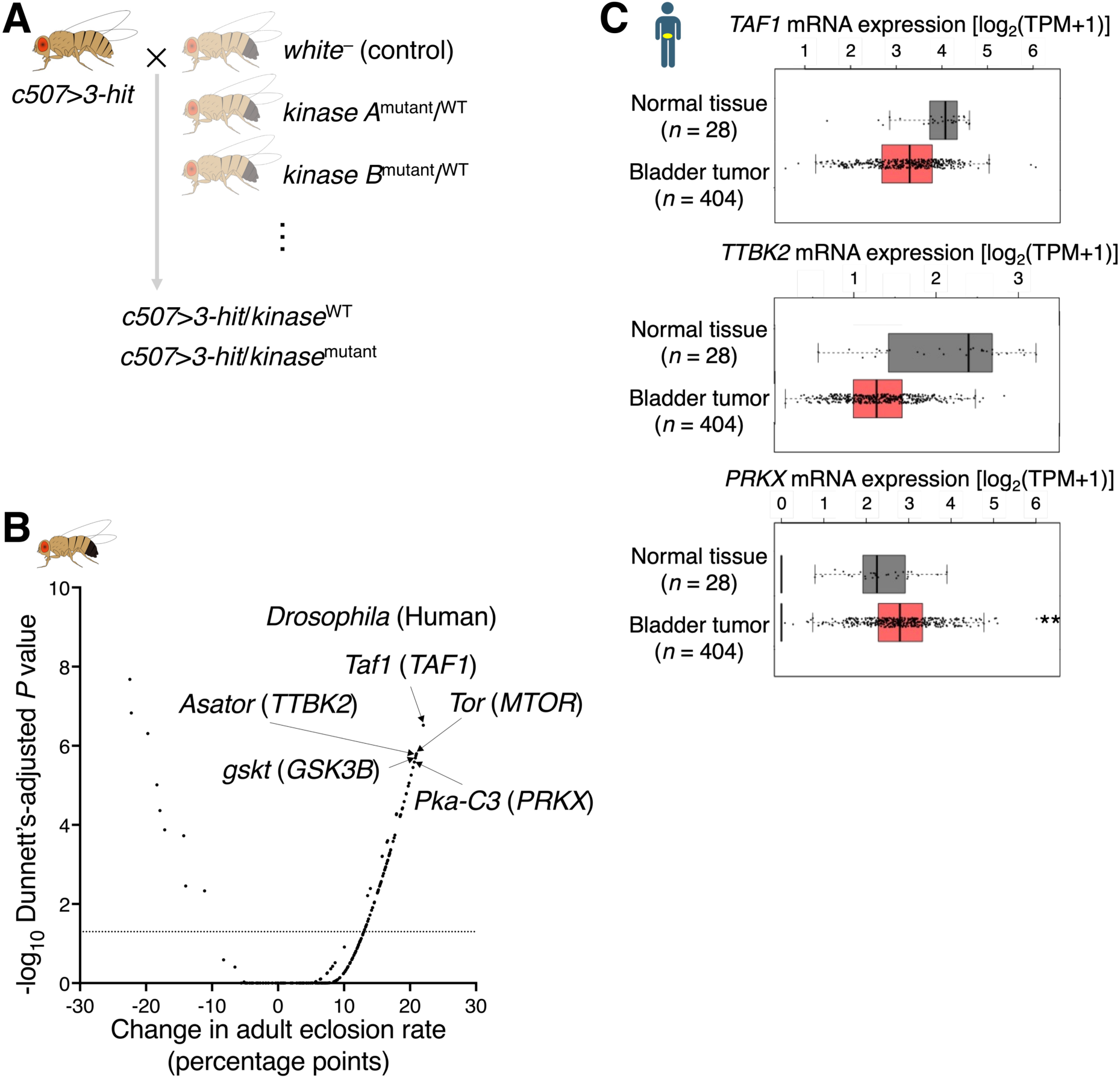
Whole-animal kinase screening identifies *Pka-C3/PRKX* as a candidate vulnerability in the *3-hit Drosophila* model. **(A)** Screening scheme for dominant modifier analysis. *c507*>*3-hit* flies were crossed with *white*− control flies or flies carrying kinase-mutant alleles. Adult eclosion rate of the resulting progeny was compared at 29°C. In the schematic, *kinase*^WT^ denotes the wild-type kinase allele and *kinase*^mutant^ denotes the mutant kinase allele. Each genotype was analyzed in three independent crosses. WT, wild type. **(B)** Kinase-screen results plotted according to the change in adult eclosion rate relative to the respective control and statistical significance. The x-axis indicates the change in adult eclosion rate in percentage points, and the y-axis indicates −log10 of the Dunnett’s-adjusted *P* value. X-linked and autosomal screens were analyzed separately against their respective controls, and the resulting values were combined only for visualization. The horizontal dashed line indicates a Dunnett’s-adjusted *P* value of 0.05. For visual clarity, the five significant suppressors showing the largest increases in adult eclosion rate are labeled with their *Drosophila* gene names and corresponding human orthologs. **(C)** GEPIA2 comparison of *TAF1*, *TTBK2*, and *PRKX* mRNA expression levels between human bladder tumors and normal bladder tissues using TCGA and GTEx data. Bladder tumors, *n* = 404; normal bladder tissues, *n* = 28. Statistical significance between tumor and normal tissues was assessed by one-way ANOVA as implemented in GEPIA2; \*\**P* < 0.01.

Mutant lines corresponding to 73 genes significantly increased the adult eclosion rate relative to their respective controls (Dunnett’s-adjusted *P* < 0.05) and were classified as suppressors of the reduced adult eclosion rate of the *3-hit* model (Fig. 2B; Fig. S3 and Table S5). Changes in adult eclosion rate and adjusted *P* values from the two screens were combined only for visualization. A complementary screen using *UAS*-kinase RNAi lines identified eight kinases whose knockdown significantly increased adult eclosion rate in the *3-hit* model (Fig. S4).

The five suppressors associated with the largest increases in adult eclosion rate were *Taf1*, *Tor*, *Asator*, *gskt*, and *Pka-C3*, corresponding to human *TAF1*, *MTOR*, *TTBK2*, *GSK3B*, and *PRKX*, respectively (Fig. 2B; Table S5). Given previous evidence implicating mTOR and GSK3β in bladder cancer cell proliferation and survival ^40–42^, we focused subsequent prioritization on *TAF1*, *TTBK2*, and *PRKX*.

We compared the expression of these three genes in bladder tumors and normal bladder tissues using GEPIA2 with TCGA and GTEx data. Both *TAF1* and *TTBK2* showed lower median expression in tumors, whereas *PRKX* expression was significantly higher (Fig. 2C). The increase in adult eclosion rate associated with *Pka-C3* perturbation, together with elevated expression of its human ortholog in tumors, supported the selection of *PRKX* for functional evaluation in human bladder cancer cells.

### Human bladder cancer cell lines show distinct *EGFR* expression and *TP53* and *RB1* mutation profiles

To extend the analysis to human bladder cancer, we selected 5637, UM-UC-3, HT-1376, and HT-1197 cells for evaluation of *PRKX*. We summarized *EGFR* expression and *TP53* and *RB1* alteration status using Cellosaurus annotations and public DepMap and CCLE datasets ^33–35^ (Table S3). *EGFR* transcript levels varied across the panel, ranging from 5.31 to 6.94 log_2_(TPM+1). Each cell line carried a different *TP53* missense variant: R280T in 5637, F113C in UM-UC-3, P250L in HT-1376, and H365R in HT-1197. Truncating *RB1* mutations were reported in 5637 (Y325*) and HT-1376 (Q702*), whereas no *RB1* alteration was reported for UM-UC-3 or HT-1197 in the referenced datasets. These profiles provided distinct molecular backgrounds for subsequent *PRKX* knockdown experiments.

### *PRKX* knockdown reduces spheroid volume across human bladder cancer models

*PRKX* is a cAMP-dependent serine/threonine kinase that is phylogenetically and functionally distinct from classical *protein kinase A* ^43^. Previous studies have implicated *PRKX* in epithelial morphogenesis, cell migration, and angiogenesis ^43–45^, and additional studies have linked *PRKX* to cancer-associated phenotypes including drug resistance and tumor-promoting signaling ^46, 47^. However, its functional role in bladder cancer remains poorly characterized.

Efficient siRNA-mediated *PRKX* knockdown was confirmed in all four bladder cancer cell lines by RT-qPCR (Fig. 3A). Representative spheroid images showed reduced spheroid size following *PRKX* knockdown (Fig. 3B). Quantitative analysis demonstrated reduced spheroid volume across all four cell lines. Both siRNAs significantly reduced spheroid volume in 5637 and UM-UC-3 cells, whereas only si*PRKX* #2 reached statistical significance in HT-1376 and HT-1197 cells (Fig. 3C). The overall pattern across cell lines with distinct *EGFR*, *TP53*, and *RB1* profiles suggests that sensitivity to *PRKX* suppression is not restricted to a single combination of the genetic features modeled in *Drosophila*. These findings support *PRKX* as a candidate vulnerability across the bladder cancer cell-line models tested.

**Figure 3.**
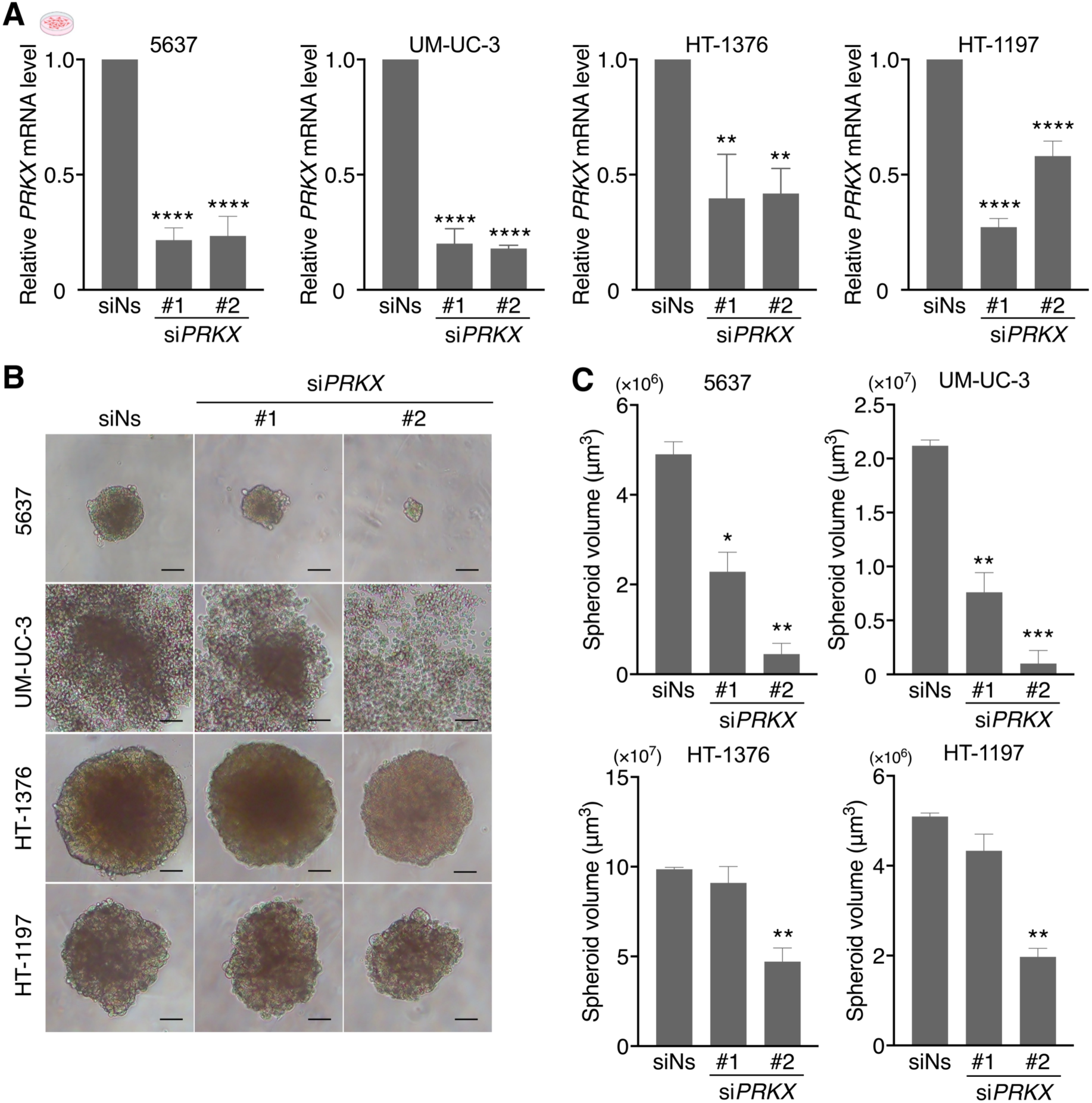
*PRKX* knockdown reduces spheroid volume across human bladder cancer models. **(A)** RT-qPCR analysis of *PRKX* mRNA levels in 5637, UM-UC-3, HT-1376, and HT-1197 cells 48 h after transfection with non-silencing control siRNA (siNs) or two independent siRNAs targeting *PRKX* (si*PRKX* #1 and #2). *PRKX* expression was normalized to *ACTB* and expressed relative to the mean expression level of the siNs group. Data are presented as mean ± SD from three independent experiments (*n* = 3). \*\**P* < 0.01 and \*\*\*\**P* < 0.0001 compared with siNs by ordinary one-way ANOVA followed by Dunnett’s multiple-comparisons test. **(B)** Representative bright-field images of spheroids formed by 5637, UM-UC-3, HT-1376, and HT-1197 cells after siRNA treatment. Images were acquired on day 7 after reseeding. Scale bars, 100 µm. **(C)** Quantification of spheroid volume. For each condition, spheroid volumes from 3–4 technical replicate wells were averaged within each independent experiment, yielding one mean value per condition per experiment. Data are presented as mean ± SD from three independent experiments per condition (*n* = 3). One spheroid was analyzed per well. Statistical significance was assessed using repeated-measures one-way ANOVA followed by Dunnett’s multiple-comparisons test, with siNs as the control group. \**P* < 0.05, \*\**P* < 0.01, and \*\*\**P* < 0.001 versus siNs.

## Discussion

In this study, we combined genetic screening in a defined *Drosophila* model with human tumor-expression analysis and functional testing in bladder cancer cells to prioritize *PRKX* as a candidate vulnerability. Perturbation of the fly ortholog *Pka-C3* increased adult eclosion rate in the *3-hit* model, and *PRKX* transcript levels were elevated in human bladder tumors. Subsequent *PRKX* knockdown reduced spheroid volume in four human bladder cancer cell lines, although the effects differed between siRNAs. This sequence of analyses connects a whole-animal genetic modifier to a phenotype in human cancer cells and provides a basis for further investigation of *PRKX* (Fig. 4).

**Figure 4.**
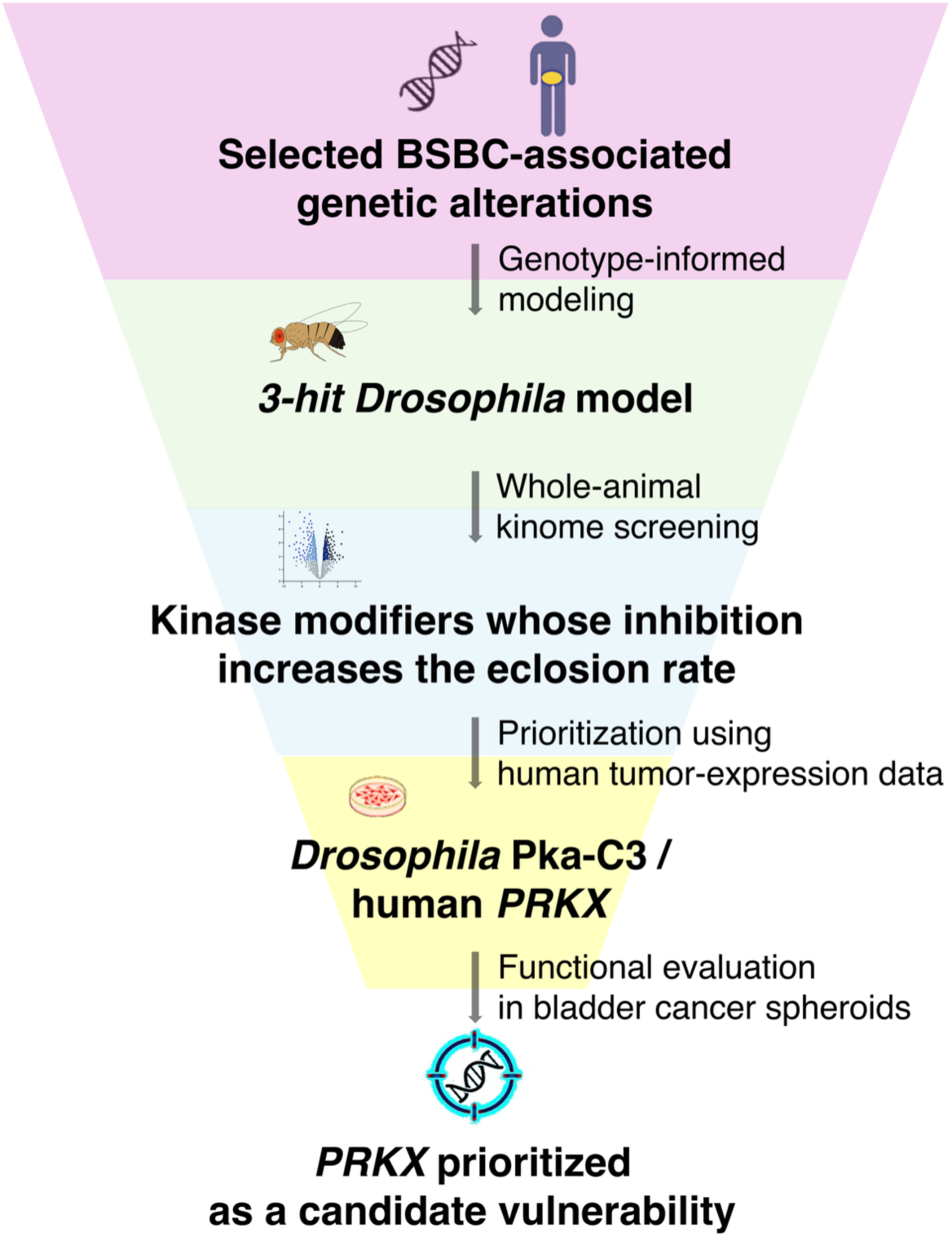
Cross-species screening prioritizes *PRKX* as a candidate bladder cancer vulnerability. Schematic overview of the cross-species candidate-prioritization strategy used in this study. A genotype-defined *Drosophila* model incorporating selected BSBC-associated genetic alterations was subjected to whole-animal kinome screening. Kinase genes whose perturbation significantly suppressed the *3-hit* phenotype were prioritized using human bladder tumor-expression data, leading to the selection of *PRKX* for functional evaluation in spheroids derived from human bladder cancer cell lines. This strategy prioritized *PRKX* as a candidate vulnerability across the models tested.

The *3-hit* model enabled systematic testing of kinase modifiers in the context of *Egfr* overexpression, mutant *p53*, and *Rbf* suppression. Expression of the *3-hit* combination in MTs expanded the GFP-positive domain and increased MT volume. In *Drosophila*, *Egfr* signaling through the *Ras–MAPK* pathway regulates cell proliferation and survival ^48^, whereas *p53* mediates cellular responses to stress, including DNA damage-induced cell death ^49, 50^. *Rbf–E2F* signaling also contributes to the control of cell-cycle progression ^51^. Against this background, the increased MT volume observed in the *3-hit* model is directionally consistent with the expected effects of simultaneously enhancing growth-promoting signaling and compromising growth-restraining pathways. However, the cellular basis of this phenotype remains unresolved. The present experiments do not distinguish whether the increased MT volume reflects increased cell proliferation, reduced cell loss, increased cell size, altered differentiation, or a combination of these processes.

This approach builds on our previous work in PDAC ^22–25^ and takes advantage of the capacity of *Drosophila* to assess genetic perturbations within a whole organism ^15, 17^. The model captures selected genetic alterations associated with BSBC ^7, 8^, rather than the broader transcriptional and microenvironmental complexity of human BSBC. Moreover, although MTs provide a tractable epithelial tissue for modeling selected cancer-associated genetic alterations ^38, 39^, they are not anatomically equivalent to the bladder. Because adult eclosion rate integrates developmental and systemic effects, increased eclosion rate in the modifier screen should be interpreted as suppression of the *3-hit* model phenotype rather than as direct evidence of reduced tumor burden. Together, these considerations define the model as a genetically controlled system for identifying modifiers of a defined BSBC-associated genetic context.

Baseline adult eclosion rate was higher in the screening crosses than in the initial characterization (Fig. 1E), which used a different crossing scheme. The parental *3-hit* stock used for screening was maintained over a *GAL80*-bearing balancer. One possible explanation is that parental carryover of *GAL80* transiently attenuated *GAL4*-dependent transgene expression during early development. However, neither maternal nor paternal transmission of functional *GAL80* was directly assessed, and the basis for the difference in baseline eclosion rate remains unresolved. Modifier effects were evaluated relative to the corresponding control within each screen.

The identification of *Tor* and *gskt*, corresponding to human *MTOR* and *GSK3B*, among the strongest suppressors is consistent with previous evidence linking these kinases to bladder cancer. Inhibition of mTOR suppresses urothelial carcinoma cell proliferation ^40, 41^, and GSK3β inhibition or depletion reduces bladder cancer cell viability ^42^. *PRKX* was selected for further study because *Pka-C3* perturbation produced one of the largest increases in adult eclosion rate and *PRKX* expression was elevated in bladder tumors. Tumor-associated expression thus complemented the genetic screen as a prioritization criterion, although it does not establish functional dependence or exclude other screen hits from further consideration.

Previous studies have linked *PRKX* to renal epithelial morphogenesis, ureteric bud branching, and endothelial proliferation, as well as to migration of renal epithelial and endothelial cells ^43–45^. In cancer models, *PRKX* has been implicated in sunitinib resistance in kidney carcinoma and melanoma cells ^46^ and in signaling associated with ovarian cancer progression ^47^. More recently, *PRKX*-mediated stabilization of PD-L1 was shown to promote immune suppression in gastric cancer ^52^. These observations suggest that *PRKX* can influence cancer-associated processes through different mechanisms. Our spheroid assays did not include immune cells, and whether *PRKX* regulates PD-L1 in bladder cancer remains unresolved.

The human cell experiments provide functional evidence beyond the tumor-expression association. Both siRNAs reduced *PRKX* mRNA levels in all four cell lines and significantly reduced spheroid volume in 5637 and UM-UC-3 cells. In HT-1376 and HT-1197 cells, only si*PRKX* #2 produced a statistically significant volume reduction. Concordant effects of both siRNAs in two cell lines, together with significant effects of si*PRKX* #2 in the other two cell lines, support an association between *PRKX* suppression and the spheroid phenotype, while the differences between siRNAs warrant further validation. Confirmation of *PRKX* protein depletion during spheroid culture, rescue with an siRNA-resistant *PRKX* construct, and independent genetic perturbation approaches would strengthen attribution of the phenotype to *PRKX* loss.

The biological basis of the spheroid-volume reduction remains to be determined. The endpoint used here cannot distinguish reduced proliferation, increased cell death, or changes in aggregation and compaction. Direct measurements of cell number, proliferation, and viability, together with time-resolved imaging, would help resolve these possibilities. Identifying *PRKX*-dependent signaling events and testing whether kinase activity is required for the phenotype would further clarify the mechanism and inform subsequent pharmacological studies.

The scope of the human validation also limits conclusions about molecular specificity. The four cell lines differed in *EGFR* expression and *TP53* and *RB1* mutation profiles, but this comparison cannot establish which features determine the response to *PRKX* suppression or whether sensitivity is enriched in BSBC. Likewise, higher *PRKX* expression in bulk bladder tumors does not establish its cellular source, protein abundance, or kinase activity. Larger panels with molecular subtype annotation, appropriate genetic comparisons, and patient-derived models will be needed to define responsive contexts. Including normal urothelial controls would also help determine whether the effects of *PRKX* suppression are selective for cancer cells.

The present findings nominate *PRKX* for mechanistic and preclinical investigation in bladder cancer. Establishing therapeutic potential will require confirmation of target specificity, evaluation in mammalian whole-body models, and assessment of pharmacological inhibition and normal-tissue tolerability. More broadly, this study illustrates how a model of selected cancer-associated alterations can guide candidate selection when whole-animal screening is followed by expression analysis and functional testing in human cells.

## Supporting information

Supplementary_Information_bioRxiv_v2

## Author contributions

**Takuya Moriguchi:** conceptualization, data curation, formal analysis, funding acquisition, validation, resources, investigation, methodology, visualization, writing – original draft, writing – review and editing. **Chihiro Kamijo:** investigation, methodology, writing – review and editing. **Taku Kimura:** resources, investigation, methodology, writing – review and editing. **Taiga Hirata:** investigation, writing – review and editing. **Reo Sato:** investigation, writing – review and editing. **Ryodai Yamamura:** funding acquisition, methodology, supervision, writing – review and editing. **Takahiro Osawa:** funding acquisition, writing – review and editing. **Nobuo Shinohara:** supervision, writing – review and editing. **Takashige Abe:** supervision, writing – review and editing. **Masahiro Sonoshita:** conceptualization, funding acquisition, project administration, resources, supervision, writing – original draft, writing – review and editing.

## Acknowledgments

We thank all members of the Sonoshita laboratory for their valuable support and discussions throughout this study. Some icons used in the figures were created in BioRender. Mo, T. (2026) https://BioRender.com/o4q73a0.

## Ethics Statement

Approval of the research protocol by an Institutional Reviewer Board: All *Drosophila* studies were conducted under protocols approved by Hokkaido University Safety Committee on Genetic Recombination Experiments (approval numbers: 2019–007 and 2022–029).

## Consent

The authors have nothing to report.

## Data Availability Statement

All data supporting the findings of this study are available from the corresponding author upon reasonable request.

## References

1 Bray F, Laversanne M, Sung H, et al. Global cancer statistics 2022: GLOBOCAN estimates of incidence and mortality worldwide for 36 cancers in 185 countries. CA Cancer J Clin. 2024; 74: 229–263. 10.3322/caac.21834

2 Powles T, Bellmunt J, Comperat E, et al. Bladder cancer: ESMO Clinical Practice Guideline for diagnosis, treatment and follow-up. Ann Oncol. 2022; 33: 244–258. 10.1016/j.annonc.2021.11.012

3 Grossman HB, Natale RB, Tangen CM, et al. Neoadjuvant chemotherapy plus cystectomy compared with cystectomy alone for locally advanced bladder cancer. N Engl J Med. 2003; 349: 859–866. 10.1056/NEJMoa022148

4 Bellmunt J, de Wit R, Vaughn DJ, et al. Pembrolizumab as Second-Line Therapy for Advanced Urothelial Carcinoma. N Engl J Med. 2017; 376: 1015–1026. 10.1056/NEJMoa1613683

5 The Cancer Genome Atlas Research Network. Comprehensive molecular characterization of urothelial bladder carcinoma. Nature. 2014; 507: 315–322. 10.1038/nature12965

6 Choi W, Porten S, Kim S, et al. Identification of distinct basal and luminal subtypes of muscle-invasive bladder cancer with different sensitivities to frontline chemotherapy. Cancer Cell. 2014; 25: 152–165. 10.1016/j.ccr.2014.01.009

7 Robertson AG, Kim J, Al-Ahmadie H, et al. Comprehensive Molecular Characterization of Muscle-Invasive Bladder Cancer. Cell. 2017; 171: 540–556.e25. 10.1016/j.cell.2017.09.007

8 Kamoun A, de Reyniès A, Allory Y, et al. A Consensus Molecular Classification of Muscle-invasive Bladder Cancer. Eur Urol. 2020; 77: 420–433. 10.1016/j.eururo.2019.09.006

9 Sjödahl G, Lauss M, Lövgren K, et al. A molecular taxonomy for urothelial carcinoma. Clin Cancer Res. 2012; 18: 3377–3386. 10.1158/1078-0432.Ccr-12-0077-t

10 Rebouissou S, Bernard-Pierrot I, de Reyniès A, et al. *EGFR* as a potential therapeutic target for a subset of muscle-invasive bladder cancers presenting a basal-like phenotype. Sci Transl Med. 2014; 6: 244ra91. 10.1126/scitranslmed.3008970

11 Brand AH, Perrimon N. Targeted gene expression as a means of altering cell fates and generating dominant phenotypes. Development. 1993; 118: 401–415. 10.1242/dev.118.2.401

12 Duffy JB. *GAL4* system in *Drosophila*: a fly geneticist’s Swiss army knife. Genesis. 2002; 34: 1–15. 10.1002/gene.10150

13 Dietzl G, Chen D, Schnorrer F, et al. A genome-wide transgenic RNAi library for conditional gene inactivation in *Drosophila*. Nature. 2007; 448: 151–156. 10.1038/nature05954

14 Pandey UB, Nichols CD. Human disease models in *Drosophila* melanogaster and the role of the fly in therapeutic drug discovery. Pharmacol Rev. 2011; 63: 411–436. 10.1124/pr.110.003293

15 Sonoshita M, Cagan RL. Modeling Human Cancers in *Drosophila*. Curr Top Dev Biol. 2017; 121: 287–309. 10.1016/bs.ctdb.2016.07.008

16 Dar AC, Das TK, Shokat KM, Cagan RL. Chemical genetic discovery of targets and anti-targets for cancer polypharmacology. Nature. 2012; 486: 80–84. 10.1038/nature11127

17 Sonoshita M, Scopton AP, Ung PMU, et al. A whole-animal platform to advance a clinical kinase inhibitor into new disease space. Nat Chem Biol. 2018; 14: 291–298. 10.1038/nchembio.2556

18 Richardson HE, Portela M. Modelling Cooperative Tumorigenesis in *Drosophila*. Biomed Res Int. 2018; 2018: 4258387. 10.1155/2018/4258387

19 Ung PMU, Sonoshita M, Scopton AP, Dar AC, Cagan RL, Schlessinger A. Integrated computational and *Drosophila* cancer model platform captures previously unappreciated chemicals perturbing a kinase network. PLoS Comput Biol. 2019; 15: e1006878. 10.1371/journal.pcbi.1006878

20 Yamamura R, Ooshio T, Sonoshita M. Tiny *Drosophila* makes giant strides in cancer research. Cancer Sci. 2021; 112: 505–514. 10.1111/cas.14747

21 Jiang H, Kimura T, Hai H, Yamamura R, Sonoshita M. *Drosophila* as a toolkit to tackle cancer and its metabolism. Front Oncol. 2022; 12: 982751. 10.3389/fonc.2022.982751

22 Sekiya S, Fukuda J, Yamamura R, et al. *Drosophila* Screening Identifies Dual Inhibition of *MEK* and *AURKB* as an Effective Therapy for Pancreatic Ductal Adenocarcinoma. Cancer Res. 2023; 83: 2704–2715. 10.1158/0008-5472.Can-22-3762

23 Fukuda J, Kosuge S, Satoh Y, et al. Concurrent targeting of *GSK3* and *MEK* as a therapeutic strategy to treat pancreatic ductal adenocarcinoma. Cancer Sci. 2024; 115: 1333–1345. 10.1111/cas.16100

24 Jiang H, Satoh Y, Yamamura R, et al. Inhibition of *NAD-GPx4* axis and *MEK* triggers ferroptosis to suppress pancreatic ductal adenocarcinoma. Mol Ther. 2025; 33: 4618– 4635. 10.1016/j.ymthe.2025.05.037

25 Hai H, Ooshio T, Jiang H, et al. Genotype-Informed Whole-Animal Kinome Screening Reveals Shared Kinase Vulnerabilities Across Driver Contexts in PDAC. Cancer Sci. 2026: 1–14. 10.1111/cas.70522

26 Bhullar KS, Lagarón NO, McGowan EM, et al. Kinase-targeted cancer therapies: progress, challenges and future directions. Mol Cancer. 2018; 17: 48. 10.1186/s12943-018-0804-2

27 Herzog G, Joerger AC, Shmueli MD, Fersht AR, Gazit E, Segal D. Evaluating *Drosophila p53* as a model system for studying cancer mutations. J Biol Chem. 2012; 287: 44330–44337. 10.1074/jbc.M112.417980

28 Bangi E, Ang C, Smibert P, et al. A personalized platform identifies trametinib plus zoledronate for a patient with *KRAS*-mutant metastatic colorectal cancer. Sci Adv. 2019; 5: eaav6528. 10.1126/sciadv.aav6528

29 Teague AG, Quintero M, Karimi Dermani F, Cagan RL, Bangi E. A polycistronic transgene design for combinatorial genetic perturbations from a single transcript in *Drosophila*. PLoS Genet. 2023; 19: e1010792. 10.1371/journal.pgen.1010792

30 McGuire SE, Le PT, Osborn AJ, Matsumoto K, Davis RL. Spatiotemporal rescue of memory dysfunction in *Drosophila*. Science. 2003; 302: 1765–1768. 10.1126/science.1089035

31 Hu Y, Comjean A, Gao C, et al. DIOPT: the DRSC Integrative Ortholog Prediction Tool, 2026 update. Genetics. 2026; 234(2): iyag194. 10.1093/genetics/iyag194

32 Tang Z, Kang B, Li C, Chen T, Zhang Z. GEPIA2: an enhanced web server for large-scale expression profiling and interactive analysis. Nucleic Acids Res. 2019; 47: W556– W560. 10.1093/nar/gkz430

33 Bairoch A. The Cellosaurus, a Cell-Line Knowledge Resource. J Biomol Tech. 2018; 29: 25–38. 10.7171/jbt.18-2902-002

34 Tsherniak A, Vazquez F, Montgomery PG, et al. Defining a Cancer Dependency Map. Cell. 2017; 170: 564–576.e16. 10.1016/j.cell.2017.06.010

35 Ghandi M, Huang FW, Jané-Valbuena J, et al. Next-generation characterization of the Cancer Cell Line Encyclopedia. Nature. 2019; 569: 503–508. 10.1038/s41586-019-1186-3

36 Livak KJ, Schmittgen TD. Analysis of relative gene expression data using real-time quantitative PCR and the 2(-Delta Delta C(T)) Method. Methods. 2001; 25: 402–408. 10.1006/meth.2001.1262

37 Kanda Y. Investigation of the freely available easy-to-use software ‘EZR’ for medical statistics. Bone Marrow Transplant. 2013; 48: 452–458. 10.1038/bmt.2012.244

38 Rodan AR. The *Drosophila* Malpighian tubule as a model for mammalian tubule function. Curr Opin Nephrol Hypertens. 2019; 28: 455–464. 10.1097/mnh.0000000000000521

39 Cohen E, Sawyer JK, Peterson NG, Dow JAT, Fox DT. Physiology, Development, and Disease Modeling in the *Drosophila* Excretory System. Genetics. 2020; 214: 235–264. 10.1534/genetics.119.302289

40 Hansel DE, Platt E, Orloff M, et al. Mammalian target of rapamycin (mTOR) regulates cellular proliferation and tumor growth in urothelial carcinoma. Am J Pathol. 2010; 176: 3062–3072. 10.2353/ajpath.2010.090872

41 Makhlin I, Zhang J, Long CJ, et al. The mTOR pathway affects proliferation and chemosensitivity of urothelial carcinoma cells and is upregulated in a subset of human bladder cancers. BJU Int. 2011; 108: E84–90. 10.1111/j.1464-410X.2010.09844.x

42 Naito S, Bilim V, Yuuki K, et al. Glycogen synthase kinase-3beta: a prognostic marker and a potential therapeutic target in human bladder cancer. Clin Cancer Res. 2010; 16: 5124–5132. 10.1158/1078-0432.Ccr-10-0275

43 Li X, Li HP, Amsler K, Hyink D, Wilson PD, Burrow CR. *PRKX*, a phylogenetically and functionally distinct cAMP-dependent protein kinase, activates renal epithelial cell migration and morphogenesis. Proc Natl Acad Sci U S A. 2002; 99: 9260–9265. 10.1073/pnas.132051799

44 Li X, Hyink DP, Polgar K, Gusella GL, Wilson PD, Burrow CR. *Protein kinase X* activates ureteric bud branching morphogenesis in developing mouse metanephric kidney. J Am Soc Nephrol. 2005; 16: 3543–3552. 10.1681/asn.2005030240

45 Li X, Iomini C, Hyink D, Wilson PD. *PRKX* critically regulates endothelial cell proliferation, migration, and vascular-like structure formation. Dev Biol. 2011; 356: 475–485. 10.1016/j.ydbio.2011.05.673

46 Bender C, Ullrich A. *PRKX*, *TTBK2* and *RSK4* expression causes Sunitinib resistance in kidney carcinoma- and melanoma-cell lines. Int J Cancer. 2012; 131: E45–55. 10.1002/ijc.26486

47 Chen Q, Xie J, Yang Y. Long non-coding RNA *NRSN2-AS1* facilitates tumorigenesis and progression of ovarian cancer via miR-744-5p/*PRKX* axis. Biol Reprod. 2022; 106: 526–539. 10.1093/biolre/ioab212

48 Yang L, Baker NE. Cell cycle withdrawal, progression, and cell survival regulation by *EGFR* and its effectors in the differentiating *Drosophila* eye. Dev Cell. 2003; 4: 359–369. 10.1016/s1534-5807(03)00059-5

49 Ollmann M, Young LM, Di Como CJ, et al. *Drosophila p53* is a structural and functional homolog of the tumor suppressor *p53*. Cell. 2000; 101: 91–101. 10.1016/s0092-8674(00)80626-1

50 Brodsky MH, Nordstrom W, Tsang G, Kwan E, Rubin GM, Abrams JM. *Drosophila p53* binds a damage response element at the reaper locus. Cell. 2000; 101: 103–113. 10.1016/s0092-8674(00)80627-3

51 Du W, Dyson N. The role of *RBF* in the introduction of G1 regulation during *Drosophila* embryogenesis. EMBO J. 1999; 18: 916–925. 10.1093/emboj/18.4.916

52 Wang Q, Chen Y, Huang D, et al. *PRKX-*mediated stabilization of PD-L1 characterizes an immunosuppressive gastric cancer subtype. J Immunother Cancer. 2026; 14: e015452. 10.1136/jitc-2026-015452

