## Supplementary_Information_bioRxiv_v2 for "Genotype-Informed *Drosophila* Kinome Screening Prioritizes *PRKX* as a Candidate Vulnerability in Bladder Cancer"

Supplementary materials and methods

Supplementary Figure S1: Schemes of fly crossing for genetic screening.

Supplementary Figure S2: The *3-hit Drosophila* model carries *Egfr* and *p53*$^{R234Q}$ transgenes and shows reduced *Rbf* expression.

Supplementary Figure S3: Complete volcano plot of the kinase screen showing changes in adult eclosion rate and adjusted *P* values for all screened kinase genes.

Supplementary Figure S4:  Complementary RNAi-based screening identifies kinase modifiers of the *3-hit Drosophila* phenotype.

Supplementary Table S1: Primer sequences used for genomic PCR and RT-qPCR.

Supplementary Table S2: List of fly strains for genetic screening.

Supplementary Table S3: Core driver alterations in human bladder cancer cell lines.

Supplementary Table S4: Culture media for human bladder cancer cell lines.

Supplementary Table S5: Significant suppressors identified in the dominant modifier screen.

### Supplementary materials and methods

#### *Fly food preparation*

Brewers' yeast (MP Biomedicals; Santa Ana, CA), yeast extract (Sigma Aldrich), bacto casitone (BD; Franklin Lakes, NJ), agar, sucrose, glucose, $MgCl_2$, $CaCl_2$, propionic acid, and mold inhibitor (10% methyl-4-hydroxybenzoate in 95% ethanol) (Fujifilm Wako; Osaka, Japan) were dissolved in water purified with reverse osmosis filters. The mixture was then boiled and cooled down.

#### *Annotation of core driver alterations in bladder cancer cell lines*

Annotations of the modeled genetic features in bladder cancer cell lines were compiled from Cellosaurus and cross-referenced with public datasets including the Cancer Dependency Map (DepMap) and Cancer Cell Line Encyclopedia (CCLE) (Supplementary Table S3).

#### *RNA interference*

*PRKX* expression was suppressed using two validated siRNAs purchased from Thermo Fisher Scientific: s11195 (si*PRKX* #1) and s224411 (si*PRKX* #2). Silencer Select Negative Control No. 1 siRNA was used as the non-silencing control (siNs). 5637, UM-UC-3, HT-1376, and HT-1197 cells were transfected with 5 nM si*PRKX* or siNs using Lipofectamine RNAiMAX (Thermo Fisher Scientific) according to the manufacturer's instructions. Knockdown efficiency was assessed by RT-qPCR 48 h after transfection.

**Figure S1. Crossing schemes for genetic screening.**
(**A**) Crossing scheme for the X-linked control screen. *3-hit* males were crossed with *white*– control females at 29°C.
(**B**) Crossing scheme for the autosomal control screen. *3-hit* females were crossed with *white*– control males at 29°C.
(**C, D**) Crossing schemes for the X-linked kinase-mutant screen. *3-hit* males were crossed with females carrying either a balanced (C) or homozygous (D) kinase-mutant allele.
(**E, F**) Crossing schemes for the 2nd chromosome kinase-mutant screen. *3-hit* females were crossed with males carrying either a balanced (E) or homozygous (F) kinase-mutant allele.
(**G, H**) Crossing schemes for the 3rd chromosome kinase-mutant screen. *3-hit* females were crossed with males carrying either a balanced (G) or homozygous (H) kinase-mutant allele.
(**I**) Crossing scheme for the 4th chromosome kinase-mutant screen. *3-hit* females were crossed with males homozygous for the kinase-mutant allele.
(**J**) Crossing scheme for the complementary kinase RNAi screen. *3-hit* females were crossed with males carrying a *UAS-kinase* RNAi transgene. Adult eclosion rate of the resulting non-*Tb* progeny was compared with that of the corresponding control cross.
All crosses were performed at 29°C. *SM5*$_{tub\text{-}GAL80}$*-TM6B* denotes an attached balancer chromosome carrying tub-GAL80, which suppresses GAL4 activity. *hs-hid* denotes the pro-apoptotic gene *hid* under the control of a heat-shock enhancer. mut, mutant.

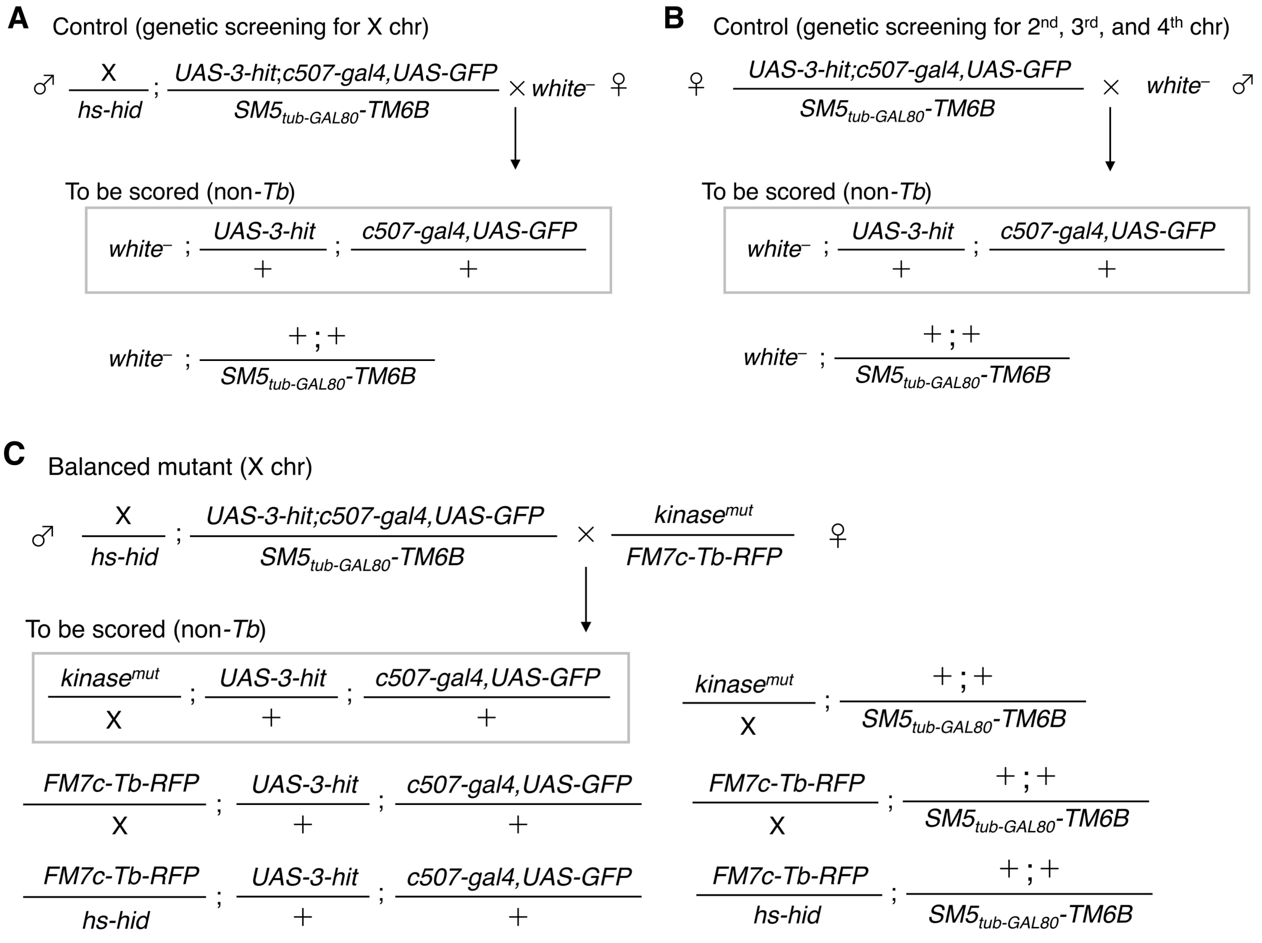

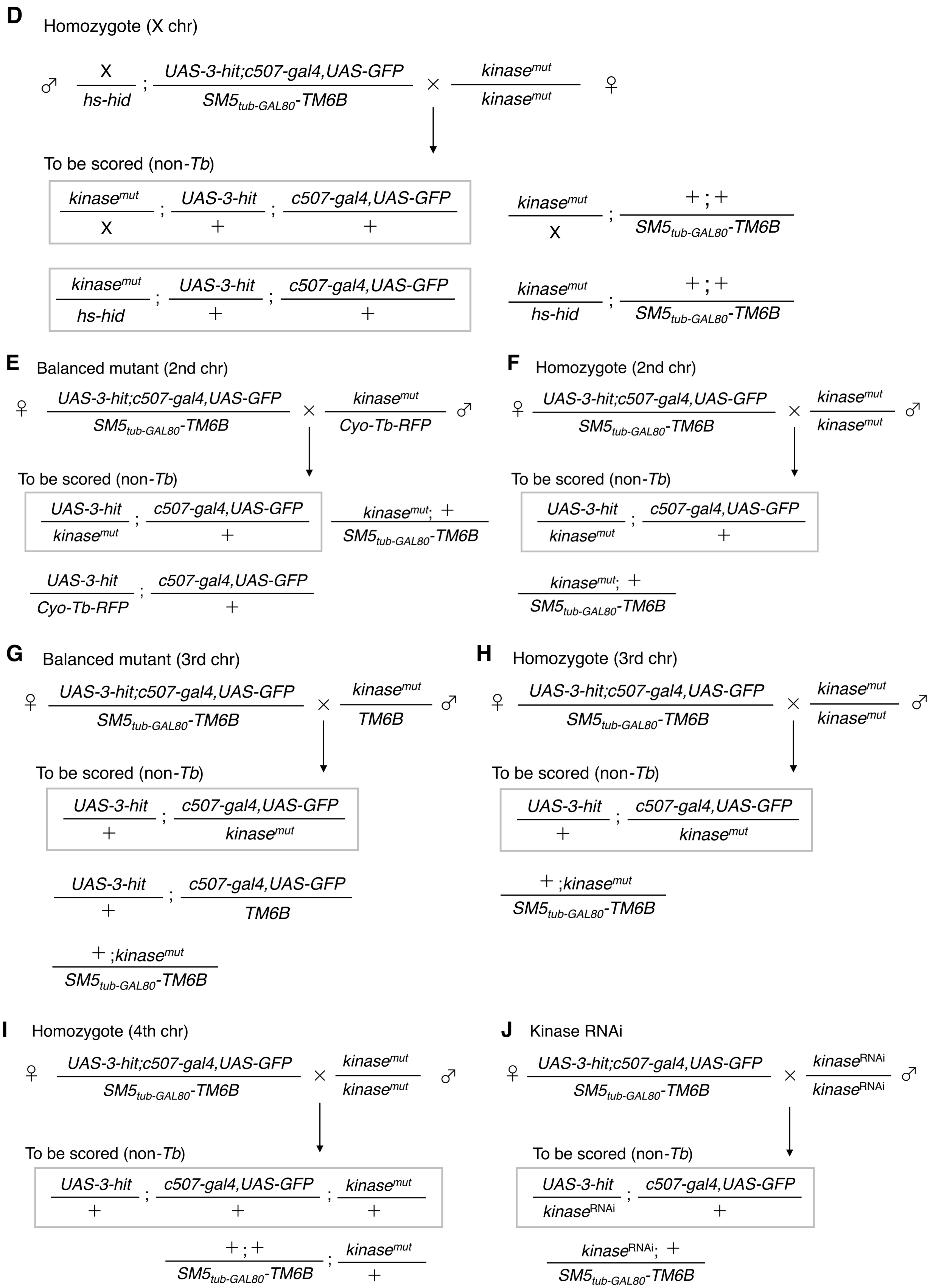
D Homozygote (X chr)
♂ X / hs-hid ; UAS-3-hit;c507-gal4,UAS-GFP / SM5tub-GAL80-TM6B × kinasemut / kinasemut ♀
To be scored (non-Tb)
kinasemut / X ; UAS-3-hit / + ; c507-gal4,UAS-GFP / +
kinasemut / X ; + ; + / SM5tub-GAL80-TM6B
kinasemut / hs-hid ; UAS-3-hit / + ; c507-gal4,UAS-GFP / +
kinasemut / hs-hid ; + ; + / SM5tub-GAL80-TM6B
E Balanced mutant (2nd chr)
♀ UAS-3-hit;c507-gal4,UAS-GFP / SM5tub-GAL80-TM6B × kinasemut / Cyo-Tb-RFP ♂
To be scored (non-Tb)
UAS-3-hit / kinasemut ; c507-gal4,UAS-GFP / +
kinasemut; + / SM5tub-GAL80-TM6B
UAS-3-hit / Cyo-Tb-RFP ; c507-gal4,UAS-GFP / +
F Homozygote (2nd chr)
♀ UAS-3-hit;c507-gal4,UAS-GFP / SM5tub-GAL80-TM6B × kinasemut / kinasemut ♂
To be scored (non-Tb)
UAS-3-hit / kinasemut ; c507-gal4,UAS-GFP / +
kinasemut; + / SM5tub-GAL80-TM6B
G Balanced mutant (3rd chr)
♀ UAS-3-hit;c507-gal4,UAS-GFP / SM5tub-GAL80-TM6B × kinasemut / TM6B ♂
To be scored (non-Tb)
UAS-3-hit / + ; c507-gal4,UAS-GFP / kinasemut
UAS-3-hit / + ; c507-gal4,UAS-GFP / TM6B
+ ;kinasemut / SM5tub-GAL80-TM6B
H Homozygote (3rd chr)
♀ UAS-3-hit;c507-gal4,UAS-GFP / SM5tub-GAL80-TM6B × kinasemut / kinasemut ♂
To be scored (non-Tb)
UAS-3-hit / + ; c507-gal4,UAS-GFP / kinasemut
+ ;kinasemut / SM5tub-GAL80-TM6B
I Homozygote (4th chr)
♀ UAS-3-hit;c507-gal4,UAS-GFP / SM5tub-GAL80-TM6B × kinasemut / kinasemut ♂
To be scored (non-Tb)
UAS-3-hit / + ; c507-gal4,UAS-GFP / + ; kinasemut / +
+ ; + / SM5tub-GAL80-TM6B ; kinasemut / +
J Kinase RNAi
♀ UAS-3-hit;c507-gal4,UAS-GFP / SM5tub-GAL80-TM6B × kinaseRNAi / kinaseRNAi ♂
To be scored (non-Tb)
UAS-3-hit / kinaseRNAi ; c507-gal4,UAS-GFP / +
kinaseRNAi; + / SM5tub-GAL80-TM6B

**Figure S2. The *3-hit Drosophila* model carries *Egfr* and *p53*$^{R234Q}$ transgenes and shows reduced *Rbf* expression.**
(**A**) Genomic PCR analysis confirming the presence of the *Egfr* and *p53*$^{R234Q}$ transgenes in *3-hit* flies. For *Egfr*, the forward primer was designed across the vector–*Egfr* insert junction, thereby distinguishing the transgene from the endogenous *Egfr* locus; the expected PCR product was 561 bp. For *p53*$^{R234Q}$, the reverse primer included sequence derived from the C-terminal c-Myc tag, enabling transgene-specific amplification; the expected PCR product was 368 bp. PCR products were separated on a 1% agarose gel in Tris-acetate-EDTA buffer. Gene Ladder Wide 1 (Nippon Gene, Tokyo, Japan) was used as the DNA size marker (M). The corresponding plasmids were used as positive controls (PCs). PCR products from *3-hit* flies are shown together with the corresponding PCs.
(**B**) RT-qPCR analysis of endogenous *Rbf* expression in Malpighian tubules from control and *3-hit* larvae. *Rbf* expression was normalized to *RpL32* and is shown relative to the control. Data are presented as mean ± SD  from three biological replicates ($n$ = 3). Statistical significance was assessed using an unpaired two-tailed Student's $t$-test; $^{**}P < 0.01$.

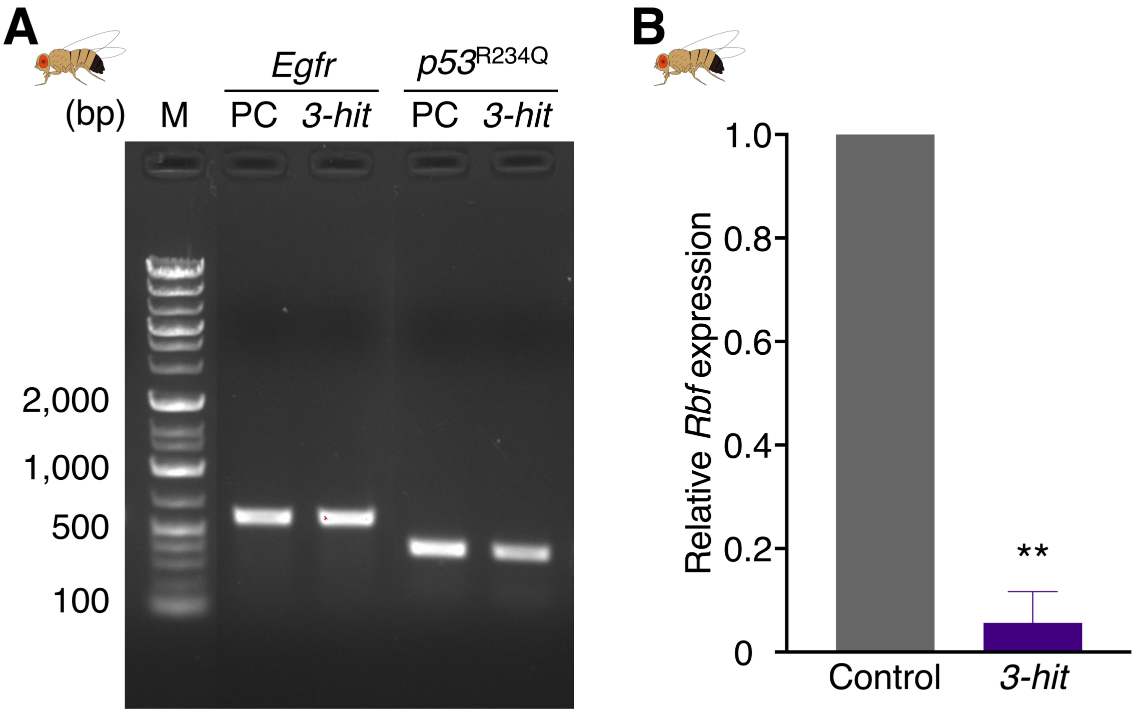

**Figure S3. Complete volcano plot of the kinase screen.**
Screening results for 216 *Drosophila* kinase genes. The x-axis indicates the change in adult eclosion rate relative to the corresponding control cross in percentage points, and the y-axis indicates −log10 of the Dunnett's-adjusted *P* value. X-linked and autosomal screens were analyzed separately against their respective control crosses before visualization in the combined plot. The horizontal dashed line indicates an adjusted *P* = 0.05.

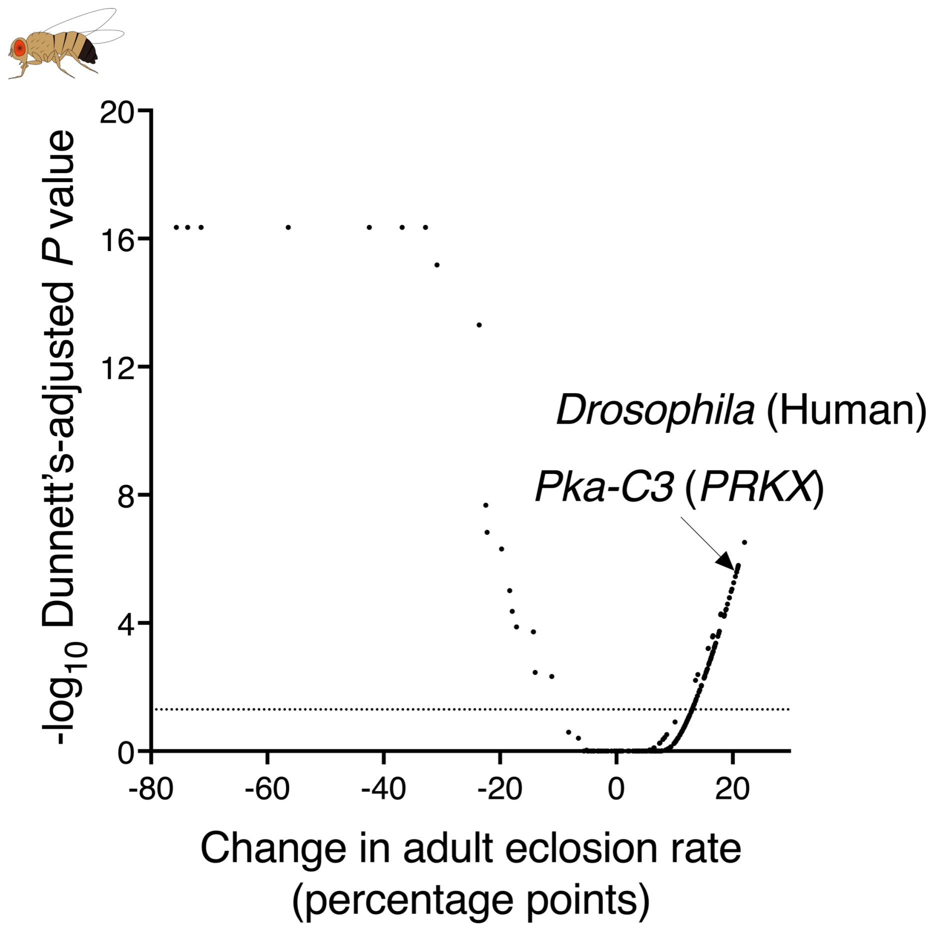

**Figure S4. Complementary RNAi-based screening identifies kinase modifiers of the *3-hit Drosophila* phenotype.**

Adult eclosion rate of *3-hit* flies crossed with the indicated kinase RNAi lines. For each genotype, adult eclosion rate was calculated as 100 × (number of eclosed non-*Tb* adults/number of non-*Tb* pupae per cross). Bars represent mean ± SD from three biological replicates (independent crosses). We assessed statistical significance relative to the control cross using one-way ANOVA with Dunnett's multiple-comparisons test (**P* < 0.05, ***P* < 0.01, ****P* < 0.001, *****P* < 0.0001).

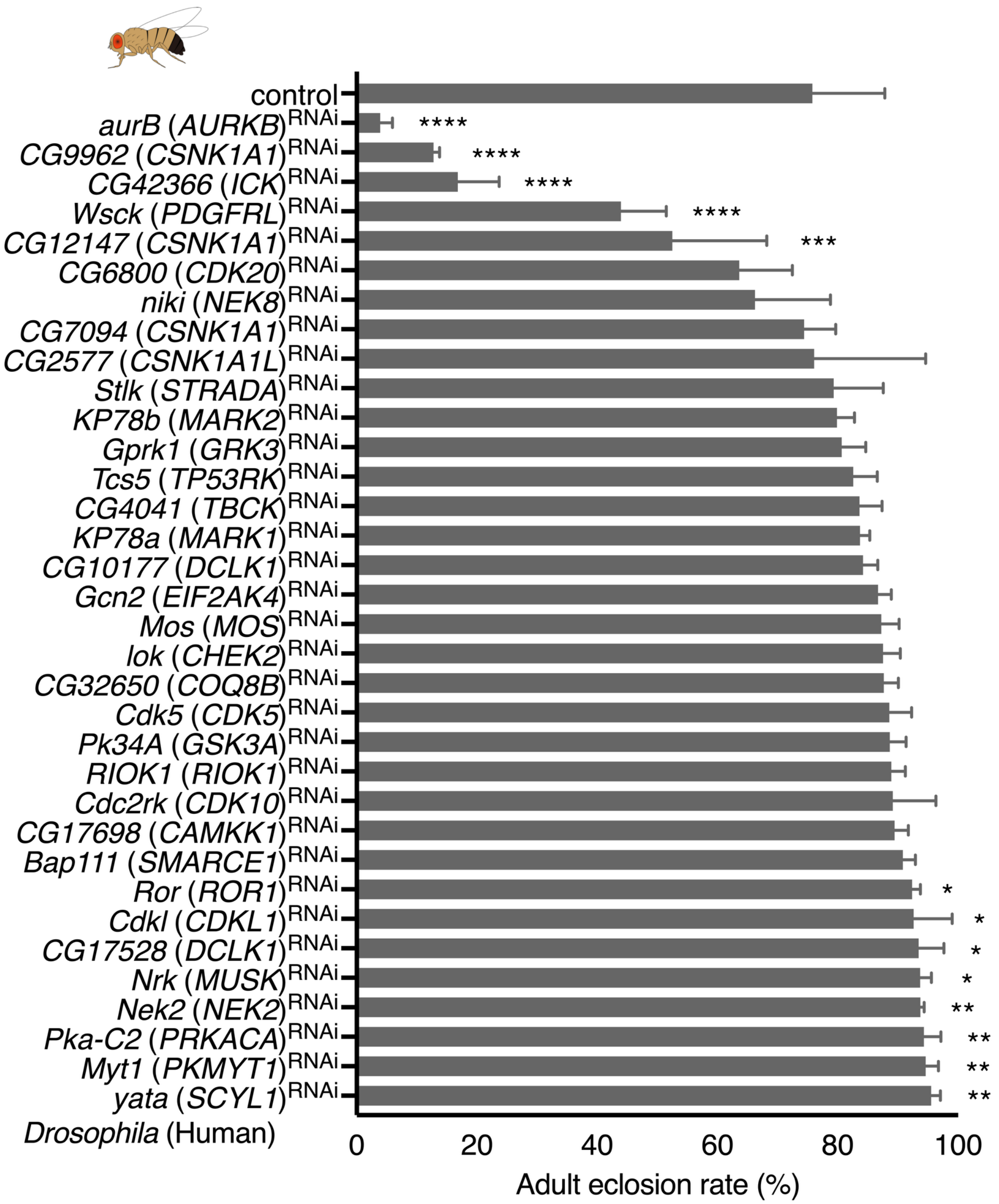

**Table S1. Primer sequences used for genomic PCR and RT-qPCR.**

| Assay | Gene | Forward | Reverse |
|---|---|---|---|
| gPCR (*Drosophila*) | *Egfr* | CTGAAGTTTCACCATGATGATTATC | GTTCGCATGTGGCAGAGAT |
| | *p53* | AGCTCAATAGCAAGAAGCG | CTCATCTCTGAAGAGGATCTGTAG |
| RT-qPCR (*Drosophila*) | *RpL32* | AGCATACAGGCCCAAGATCG | TGTTGTCGATACCCTTGGGC |
| | *Rbf* | ACACAAACCACACACGCCCAAATC | CTAGTGGCTGCCGGTGTATG |
| RT-qPCR (BC cell lines) | *ACTB* | AGAAAATCTGGCACCACACC | TGGGGTGTTGAAGGTCTCA |
| | *PRKX* | CTGGACGTGGCATGACGAG | TCGATGGCACAGATGATCTCT |

**Table S2. List of fly strains for genetic screening.**

| CG No. | Fly gene | Human ortholog | Allele / RNAi transgene | Stock center | Stock No. | RRID |
|---|---|---|---|---|---|---|
| CG4032 | *Abl* | *ABL* | *Abl*[2] | BDSC | 8565 | BDSC_8565 |
| CG14992 | *Ack* | *TNK2* | P{EPgy2}*Ack*[EY09374] | BDSC | 22316 | BDSC_22316 |
| CG43741 | *Ack-like* | *TNK1* | Mi{ET1}*Ack-like*[MB05119] | BDSC | 24322 | BDSC_24322 |
| CG3608 | *Adck1* | *ADCK1* | PBac{WH}*Adck* [f03261] | BDSC | 18633 | BDSC_18633 |
| CG7616 | *Adck5* | *ADCK5* | P{EP}*Adck5*[G14668] | BDSC | 28404 | BDSC_28404 |
| CG8866 | *Aduk* | *ULK3* | P{EPgy2}*Aduk*[EY18321] | BDSC | 17323 | BDSC_17323 |
| CG4006 | *Akt1* | *AKT1* | P{EPgy2}*Akt*[EY10012] | BDSC | 19894 | BDSC_19894 |
| CG8250 | *Alk* | *ALK* | Mi{ET1}*Alk*[MB06458] | BDSC | 25509 | BDSC_25509 |
| CG3051 | *AMPKalpha* | *PRKAA2* | *AMPKalpha*[1] | BDSC | 33830 | BDSC_33830 |
| CG42783 | *aPKC* | *PRKCI* | P{lacW}*aPKC*[k06403] | BDSC | 10622 | BDSC_10622 |
| CG11533 | *Asator* | *TTBK2* | P{SUPor-P}*Asator*[KG05051] | BDSC | 13625 | BDSC_13625 |
| CG4720 | *Ask1* | *MAP3K15* | Mi{MIC}*Ask1*[MI02915] | BDSC | 36163 | BDSC_36163 |
| CG10967 | *Atg1* | *ULK1* | *Atg1*[00305] | BDSC | 11494 | BDSC_11494 |
| CG3068 | *aurA* | *AURKA* | *aurA*[87Ac-3] | BDSC | 6188 | BDSC_6188 |
| CG6620 | *aurB* | *AURKB* | P{TRiP.JF03107}attP2 (*aurB* RNAi) | BDSC | 28691 | BDSC_28691 |
| CG1107 | *aux* | *GAK* | *aux*[L7] | BDSC | 25674 | BDSC_25674 |
| CG8224 | *babo* | *TGFBR1* | *babo*[32] | BDSC | 5399 | BDSC_5399 |
| CG6386 | *ball* | *VRK1* | Mi{MIC}*ball*[MI14724] | BDSC | 59723 | BDSC_59723 |
| CG7055 | *Bap111* | *SMARCE1* | P{TRiP.GL00129}attP2 (*Bap111* RNAi) | BDSC | 35242 | BDSC_35242 |
| CG34356 | *bma* | *SCYL2* | Mi{MIC}*bma*[MI08649] | BDSC | 51097 | BDSC_51097 |
| CG5206 | *bon* | *TRIM24* | P{lacW}*bon*[S048706] | BDSC | 4543 | BDSC_4543 |
| CG5680 | *bsk* | *JNK* | *bsk*[1] | BDSC | 3088 | BDSC_3088 |
| CG32019 | *bt* | *TTN* | *bt*[l-b] | BDSC | 4231 | BDSC_4231 |
| CG8049 | *Btk29A* | *TEC* | P{lacW}*Btk*[k00206] | BDSC | 10469 | BDSC_10469 |
| CG32134 | *btl* | *FGFR* | P{EPgy2}*btl*[EY01638] | BDSC | 15076 | BDSC_15076 |
| CG14030 | *Bub1* | *BUB1* | PBac{PB}*Bub1*[c04512] | BDSC | 11489 | BDSC_11489 |
| CG7838 | *BubR1* | *BUB1B* | P{lacW}*BubR1*[k03113] | BDSC | 10526 | BDSC_10526 |
| CG10244 | *Cad96Ca* | *KDR* | P{XP}*Cad96Ca*[d07355] | BDSC | 19270 | BDSC_19270 |
| CG1495 | *CaMKI* | *CAMK1D* | P{EPgy2}*CaMKI*[EY07197] | BDSC | 16799 | BDSC_16799 |
| CG18069 | *CaMKII* | *CAMK2D* | P{EPgy2}*CaMKII*[EY14097] | BDSC | 22325 | BDSC_22325 |
| CG6703 | *CASK* | *CASK* | *CASK*[X-313] | BDSC | 6784 | BDSC_6784 |
| CG1362 | *Cdc2rk* | *CDK10* | P{TRiP.GL00232}attP2 (*Cdc2rk* RNAi) | BDSC | 35325 | BDSC_35325 |
| CG32742 | *Cdc7* | *CDC7* | P{lacW}*Cdc7*[G0148] | BDSC | 11937 | BDSC_11937 |
| CG6027 | *cdi* | *TESK2* | *cdi*[R47] | BDSC | 9587 | BDSC_9587 |
| CG5363 | *Cdk1* | *CDK1* | *Cdk1*[E1-23] | BDSC | 6629 | BDSC_6629 |
| CG7597 | *Cdk12* | *CDK12* | P{SUPor-P}*Cdk12*[KG05512] | BDSC | 14118 | BDSC_14118 |
| CG10498 | *Cdk2* | *CDK2* | *Cdk2*[2] | BDSC | 6632 | BDSC_6632 |
| CG5072 | *Cdk4* | *CDK4/6* | P{lacW}*Cdk4*[s4639] | BDSC | 12056 | BDSC_12056 |
| CG8203 | *Cdk5* | *CDK5* | P{TRiP.JF02667}attP2 (*Cdk5* RNAi) | BDSC | 27517 | BDSC_27517 |
| CG3319 | *Cdk7* | *CDK7* | *Cdk7*[A] | BDSC | 57067 | BDSC_57067 |
| CG10572 | *Cdk8* | *CDK8* | *Cdk8*[A162] | BDSC | 16058 | BDSC_16058 |

| | | | | | | |
|---|---|---|---|---|---|---|
| CG5179 | *Cdk9* | *CDK9* | PBac{WH}*Cdk9*[f05537] | BDSC | 18881 | BDSC_18881 |
| CG7236 | *Cdkl* | *CDKL1* | P{TRiP.JF02655}attP2 (*Cdkl* RNAi) | BDSC | 27505 | BDSC_27505 |
| CG10177 | *CG10177* | *DCLK1* | P{TRiP.JF01965}attP2 (*CG10177* RNAi) | BDSC | 25945 | BDSC_25945 |
| CG12069 | *CG12069* | *PRKACA* | Mi{ET1}*CG12069*[MB10013] | BDSC | 27824 | BDSC_27824 |
| CG12147 | *CG12147* | *CSNK1A1* | P{TRiP.HMS02447}attP2 (*CG12147* RNAi) | BDSC | 42612 | BDSC_42612 |
| CG1227 | *CG1227* | *STK16* | PBac{WH}*CG1227*[f02638] | BDSC | 85651 | BDSC_85651 |
| CG1344 | *CG1344* | *SCYL3* | P{EP}*CG1344*[EP2646] | BDSC | 17079 | BDSC_17079 |
| CG17528 | *CG17528* | *DCLK1* | P{TRiP.JF02061}attP2 (*CG17528* RNAi) | BDSC | 26292 | BDSC_26292 |
| CG17698 | *CG17698* | *CAMKK1* | P{TRiP.HMS01475}attP2 (*CG17698* RNAi) | BDSC | 35733 | BDSC_35733 |
| CG1951 | *CG1951* | *SCYL2* | P{EPgy2}*CG1951*[EY00129] | BDSC | 14809 | BDSC_14809 |
| CG2577 | *CG2577* | *CSNK1A1L* | P{TRiP.HMS00054}attP2 (*CG2577* RNAi) | BDSC | 33649 | BDSC_33649 |
| CG3008 | *CG3008* | *RIOK3* | P{EP}*CG3008*[G18059] | BDSC | 27428 | BDSC_27428 |
| CG31183 | *CG31183* | *NPR2* | Mi{MIC}*CG31183*[MI02001] | BDSC | 33160 | BDSC_33160 |
| CG3216 | *CG3216* | *NPR1* | Mi{ET1}*CG3216*[MB07455] | BDSC | 26080 | BDSC_26080 |
| CG32650 | *CG32650* | *COQ8B* | UAS-*CG32650* RNAi | VDRC | 109827 | not available |
| CG3277 | *CG3277* | *STYK1* | Mi{MIC}*CG3277*[MI06697] | BDSC | 43078 | BDSC_43078 |
| CG4041 | *CG4041* | *TBCK* | P{TRiP.GL00239}attP2 (*CG4041* RNAi) | BDSC | 35332 | BDSC_35332 |
| CG42366 | *CG42366* | *ICK* | P{TRiP.JF02599}attP2 (CG42366 RNAi) | BDSC | 27294 | BDSC_27294 |
| CG42637 | *CG42637* | *NPR1* | Mi{MIC}*CG42637*[MI10816] | BDSC | 55544 | BDSC_55544 |
| CG4629 | *CG4629* | *NIM1* | Mi{MIC}*CG4629*[MI02585] | BDSC | 36008 | BDSC_36008 |
| CG4839 | *CG4839* | *PRKG1* | Mi{MIC}*CG4839*[MI08552] | BDSC | 44791 | BDSC_44791 |
| CG4945 | *CG4945* | *SBK2* | *CG4945*[f02115] | BDSC | 18515 | BDSC_18515 |
| CG5790 | *CG5790* | *CDC7* | PBac{WH}*CG5790*[f04763] | BDSC | 18803 | BDSC_18803 |
| CG6800 | *CG6800* | *CDK20* | P{TRiP.HMC05159}attP40 (*CG6800* RNAi) | BDSC | 62152 | BDSC_62152 |
| CG7094 | *CG7094* | *CSNK1A1* | P{TRiP.HMS02489}attP2 (*CG7094* RNAi) | BDSC | 42653 | BDSC_42653 |
| CG8173 | *CG8173* | *PBK* | P{RS3}*CG8173*[CB-5632-3] | DGRC | 123720 | not available |
| CG8565 | *CG8565* | *SRPK3* | Mi{MIC}*CG8565*[MI12034] | BDSC | 56849 | BDSC_56849 |
| CG8726 | *CG8726* | *PXK* | P{EPgy2}*CG8726*[EY21837] | BDSC | 22513 | BDSC_22513 |
| CG8878 | *CG8878* | *VRK3* | P{EPgy2}*CG8878*[EY10775] | BDSC | 20213 | BDSC_20213 |
| CG9962 | *CG9962* | *CSNK1A1* | P{TRiP.HMC05154}attP40 (*CG9962* RNAi) | BDSC | 62148 | BDSC_62148 |
| CG2028 | *CkIalpha* | *CSNK1A1* | P{lacW}*CkIalpha*[G0492] | BDSC | 12303 | BDSC_12303 |
| CG17520 | *CkIIalpha* | *CSNK2A1* | *CkIIalpha*[Tik] | BDSC | 24512 | BDSC_24512 |
| CG32649 | *Coq8* | *COQ8A* | *Coq8*[A] | BDSC | 52331 | BDSC_52331 |
| CG42317 | *Csk* | *CSK* | *Csk*[j1D8] | BDSC | 10244 | BDSC_10244 |
| CG2048 | *dco* | *CSNK1E* | P{lacW}*dco*[j3B9] | BDSC | 12164 | BDSC_12164 |
| CG33531 | *Ddr* | *DDR2* | Mi{y[+mDint2]=MIC}*Ddr*[MI04117] | BDSC | 37241 | BDSC_37241 |
| CG8657 | *Dgkepsilon* | *DGKE* | *Dgkepsilon*[ox-1] | BDSC | 11754 | BDSC_11754 |
| CG17559 | *dnt* | *RYK* | PBac{PB}*dnt*[c03766] | BDSC | 85366 | BDSC_85366 |
| CG3727 | *dock* | *NCK1* | P{PZ}*dock*[04723] | BDSC | 11385 | BDSC_11385 |
| CG6498 | *dop* | *MAST1* | Mi{ET1}*dop*[MB04862] | BDSC | 24830 | BDSC_24830 |
| CG32666 | *Drak* | *STK17A, STK17B* | P{SUPor-P}*Drak*[KG03058] | BDSC | 13768 | BDSC_13768 |
| CG17348 | *drl* | *RYK* | *drl*[exc21] | BDSC | 7363 | BDSC_7363 |
| CG3915 | *Drl-2* | *RYK* | P{GT1}*Drl-2*[BG02105] | BDSC | 12655 | BDSC_12655 |
| CG15793 | *Dsor1* | *MEK* | *Dsor1*[LH110] | BDSC | 5545 | BDSC_5545 |

| | | | | | | |
|---|---|---|---|---|---|---|
| CG4551 | *Dyrk2* | *DYRK4* | Mi{MIC}*Dyrk2*[MI04771] | BDSC | 38165 | BDSC_38165 |
| CG40478 | *Dyrk3* | *DYRK3* | Mi{MIC}*Dyrk3*[MI06694] | BDSC | 60800 | BDSC_60800 |
| CG3525 | *eas* | *ETNK1* | *eas*[alaE13] | BDSC | 9463 | BDSC_9463 |
| CG10079 | *Egfr* | *EGFR* | *Egfr*[f2] | BDSC | 2768 | BDSC_2768 |
| CG10738 | *EHR* | *NPR1* | Mi{ET1}*EHR*[MB04068] | BDSC | 24564 | BDSC_24564 |
| CG10579 | *Eip63E* | *CDK14* | *Eip63E*[81] | BDSC | 4513 | BDSC_4513 |
| CG1511 | *Eph* | *EPHB1* | Mi{MIC}*Eph*[MI05205] | BDSC | 61713 | BDSC_61713 |
| CG32703 | *Erk7* | *MAPK15* | Mi{MIC}*Erk7*[MI05843] | BDSC | 43776 | BDSC_43776 |
| CG10023 | *Fak* | *PTK2* | P{SUPor-P}*Fak*[KG00304] | BDSC | 13080 | BDSC_13080 |
| CG8874 | *FER* | *FER* | P{lacW}*FER*[X42] | BDSC | 9361 | BDSC_9361 |
| CG10033 | *for* | *PRKG1* | P{UASp-YFP.RabX2.S21N}*for*[02] | BDSC | 9843 | BDSC_9843 |
| CG7693 | *fray* | *STK39* | Mi{MIC}*fray*[MI03454] | BDSC | 37193 | BDSC_37193 |
| CG2252 | *fs(1)h* | *BRD2* | *fs(1)h*[rnc] | BDSC | 5278 | BDSC_5278 |
| CG6551 | *fu* | *STK36* | *fu*[mH63] | BDSC | 7207 | BDSC_7207 |
| CG5169 | *GckIII* | *MST4* | P{EPgy2}*GckIII*[EY05076] | BDSC | 15946 | BDSC_15946 |
| CG1609 | *Gcn2* | *EIF2AK4* | P{TRiP.GL00267}attP2 (*Gcn2* RNAi) | BDSC | 35355 | BDSC_35355 |
| CG4012 | *gek* | *CDC42BPA* | P{PZ}*gek*[09373] | BDSC | 12361 | BDSC_12361 |
| CG6963 | *gish* | *CSNK1G3* | P{SUPor-P}*gish*[KG03891] | BDSC | 13263 | BDSC_13263 |
| CG40129 | *Gprk1* | *GRK3* | P{TRiP.JF02990}attP2 (*Gprk1* RNAi) | BDSC | 28354 | BDSC_28354 |
| CG17998 | *Gprk2* | *GRK5* | P{PZ}*Gprk2*[06936] | BDSC | 10351 | BDSC_10351 |
| CG17161 | *grp* | *CHEK1* | P{PZ}*grp*[06034] | BDSC | 12219 | BDSC_12219 |
| CG31003 | *gskt* | *GSK3B* | Mi{MIC}*gskt*[MI04964] | BDSC | 37883 | BDSC_37883 |
| CG34357 | *Gucy2d* | *GUCY2D* | P{EPgy2}*Gucy2d*[EY21024] | BDSC | 22444 | BDSC_22444 |
| CG7719 | *gwl* | *MASTL* | P{EP}*gwl*[EP515] | BDSC | 17174 | BDSC_17174 |
| CG33114 | *Gyc32E* | *NPR2* | P{SUPor-P}*Gyc32E*[KG06014] | BDSC | 14137 | BDSC_14137 |
| CG42636 | *Gyc76C* | *NPR1* | *Gyc76C*[A377] | BDSC | 16101 | BDSC_16101 |
| CG40080 | *Haspin* | *GSG2* | Mi{MIC}*Haspin*[MI09386] | BDSC | 53099 | BDSC_53099 |
| CG4353 | *hep* | *MAP2K7* | *hep*[r75] | BDSC | 6761 | BDSC_6761 |
| CG17090 | *hipk* | *HIPK2* | Mi{MIC}*Hipk*[MI11849] | BDSC | 56388 | BDSC_56388 |
| CG1594 | *hop* | *JAK* | *hop*[2] | BDSC | 6032 | BDSC_6032 |
| CG11228 | *hpo* | *STK3* | *hpo*[KC202] | BDSC | 25090 | BDSC_25090 |
| CG7097 | *hppy* | *MAP4K3* | P{lacW}*hppy*[SH1261] | BDSC | 29493 | BDSC_29493 |
| CG7223 | *htl* | *FGFR* | *htl*[AB42] | BDSC | 5370 | BDSC_5370 |
| CG4201 | *IKKbeta* | *CHUK* | *IKKbeta*[1] | BDSC | 30916 | BDSC_30916 |
| CG2615 | *IKKepsilon* | *TBK1* | *IKKepsilon*[5] | BDSC | 5324 | BDSC_5324 |
| CG10504 | *Ilk* | *ILK* | *Ilk*[1] | BDSC | 4861 | BDSC_4861 |
| CG6518 | *inaC* | *PRKCA* | *inaC*[2] | BDSC | 42241 | BDSC_42241 |
| CG18402 | *InR* | *INSR* | P{PZ}*InR*[05545] | BDSC | 11661 | BDSC_11661 |
| CG4583 | *Ire1* | *ERN1* | PBac{WH}*Ire1*[f02170] | BDSC | 18520 | BDSC_18520 |
| CG6297 | *JIL-1* | *RPS6KA5* | P{SUPor-P}*JIL-1*[KG02848] | BDSC | 13330 | BDSC_13330 |
| CG6715 | *KP78a* | *MARK1* | UAS-*KP78a* RNAi | VDRC | 26722 | not available |
| CG17216 | *KP78b* | *MARK2* | UAS-*KP78b* RNAi | VDRC | 105265 | not available |
| CG2899 | *ksr* | *KSR* | P{EPgy2}*ksr*[EY01688] | BDSC | 19645 | BDSC_19645 |
| CG12244 | *lic* | *MAP2K6* | P{Mae-UAS.6.11}*lic*[GG01785] | BDSC | 19989 | BDSC_19989 |

| | | | | | | |
|---|---|---|---|---|---|---|
| CG1848 | *LIMK1* | *LIMK1* | P{EPgy2}*LIMK1*[EY08757] | BDSC | 17491 | BDSC_17491 |
| CG17342 | *Lk6* | *MKNK1* | *Lk6*[2] | BDSC | 8707 | BDSC_8707 |
| CG9374 | *Lkb1* | *LKB1* | P{EP}*Lkb1*[G5285] | BDSC | 30148 | BDSC_30148 |
| CG10895 | *lok* | *CHEK2* | P{TRiP.HMC05499}attP40 (*lok* RNAi) | BDSC | 64482 | BDSC_64482 |
| CG5483 | *Lrrk* | *LRRK1* | *Lrrk*[ex1] | BDSC | 34750 | BDSC_34750 |
| CG1098 | *Madm* | *NRBP1* | P{EP}*Madm*[EP3137] | BDSC | 17093 | BDSC_17093 |
| CG3086 | *MAPk-Ak2* | *MAPKAPK3* | P{EP}*MAPk-Ak2*[G265] | BDSC | 33524 | BDSC_33524 |
| CG18582 | *mbt* | *PAK7* | P{EPgy2}*mbt*[EY08341] | BDSC | 17453 | BDSC_17453 |
| CG4252 | *mei-41* | *ATR* | *mei-41*[RT1] | BDSC | 4169 | BDSC_4169 |
| CG7717 | *Mekk1* | *MAP3K4* | P{EPgy2}*Mekk1*[EY11461] | BDSC | 20676 | BDSC_20676 |
| CG11221 | *meng* | *SBK1* | Mi{MIC}*meng*[MI03008] | BDSC | 36177 | BDSC_36177 |
| CG9738 | *Mkk4* | *MAP2K4* | PBac{RB}*Mkk4*[e01485] | BDSC | 17956 | BDSC_17956 |
| CG42273 | *mnb* | *DYRK1B* | P{EPgy2}*mnb*[EY14320] | BDSC | 21430 | BDSC_21430 |
| CG8767 | *Mos* | *MOS* | P{TRiP.GL00142}attP2 (*Mos* RNAi) | BDSC | 35254 | BDSC_35254 |
| CG7643 | *Mps1* | *TTK* | P{GSV1}*Mps1*[EP-M50.2] | BDSC | 43462 | BDSC_43462 |
| CG16973 | *msn* | *MAP4K4* | *msn*[102] | BDSC | 5945 | BDSC_5945 |
| CG32417 | *Myt1* | *PKMYT1* | P{TRiP.GL00216}attP2 (*Myt1* RNAi) | BDSC | 35312 | BDSC_35312 |
| CG10637 | *Nak* | *AAK1* | PBac{WH}*Nak*[f04720] | BDSC | 18799 | BDSC_18799 |
| CG6343 | *ND-42* | *NDUFA10* | P{EP}*ND-42*[G4500] | BDSC | 30085 | BDSC_30085 |
| CG17256 | *Nek2* | *NEK2* | P{TRiP.GL00235}attP2 (*Nek2* RNAi) | BDSC | 35328 | BDSC_35328 |
| CG10951 | *niki* | *NEK8* | P{TRiP.HMS01477}attP2 (*niki* RNAi) | BDSC | 35735 | BDSC_35735 |
| CG5125 | *ninaC* | *MYO3A* | *ninaC*[5] | BDSC | 1996 | BDSC_1996 |
| CG33554 | *Nipped-A* | *TRRAP* | *Nipped-A*[NC186] | BDSC | 7187 | BDSC_7187 |
| CG7892 | *nmo* | *NLK* | *nmo*[j147-1] | BDSC | 5407 | BDSC_5407 |
| CG32743 | *nonC* | *SMG1* | P{EP}*nonC*[G1076] | BDSC | 26614 | BDSC_26614 |
| CG4007 | *Nrk* | *MUSK* | P{EP}*Nrk*[G2759] | BDSC | 28454 | BDSC_28454 |
| CG4007 | *Nrk* | *MUSK* | P{TRiP.HMC04375}attP2 (*Nrk* RNAi) | BDSC | 56936 | BDSC_56936 |
| CG43143 | *Nuak* | *NUAK1* | Mi{ET1}*Nuak*[MB02272] | BDSC | 23429 | BDSC_23429 |
| CG33519 | *Obsc* | *SPEG* | P{EPgy2}*Obsc*[EY15484] | BDSC | 21139 | BDSC_21139 |
| CG8967 | *otk* | *PTK7* | P{EP}*otk*[EP2017] | BDSC | 17231 | BDSC_17231 |
| CG5475 | *p38a* | *MAPK14* | *p38a*[1] | BDSC | 8822 | BDSC_8822 |
| CG7393 | *p38b* | *MAPK14* | P{EPgy2}*p38b*[EY11174] | BDSC | 22319 | BDSC_22319 |
| CG33338 | *p38c* | *MAPK11* | P{SUPor-P}*p38c*[KG05834] | BDSC | 14126 | BDSC_14126 |
| CG10295 | *Pak* | *PAK1* | *Pak*[6] | BDSC | 8809 | BDSC_8809 |
| CG14895 | *Pak3* | *PAK3* | P{Mae-UAS.6.11}*Pak3*[LA00012] | BDSC | 22211 | BDSC_22211 |
| CG8201 | *par-1* | *MARK3* | P{lacW}*par-1*[k06323] | BDSC | 10615 | BDSC_10615 |
| CG3105 | *Pask* | *PASK* | Mi{MIC}*Pask*[MI04252] | BDSC | 37625 | BDSC_37625 |
| CG8808 | *Pdk* | *PDK3* | P{EPgy2}*Pdk*[EY01879] | BDSC | 15084 | BDSC_15084 |
| CG1210 | *Pdk1* | *PDPK1* | P{SUPor-P}*Pdk1*[KG01447] | BDSC | 13441 | BDSC_13441 |
| CG2087 | *PEK* | *EIF2AK3* | P{EPgy2}*PEK*[EY09578] | BDSC | 17582 | BDSC_17582 |
| CG1830 | *PhKγ* | *PHKG2* | P{EPgy2}*PhKgamma*[EY12008] | BDSC | 20720 | BDSC_20720 |
| CG11621 | *Pi3K68D* | *PIK3C2A* | Mi{MIC}*Pi3K68D*[MI15179] | BDSC | 59605 | BDSC_59605 |
| CG4141 | *Pi3K92E* | *PIK3CA* | Mi{ET1}*Pi3K92E*[MB06212] | BDSC | 25483 | BDSC_25483 |
| CG4523 | *Pink1* | *PINK1* | *Pink1*[B9] | BDSC | 34749 | BDSC_34749 |

| | | | | | | |
|---|---|---|---|---|---|---|
| CG4268 | *Pitslre* | *CDK11B* | P{EPgy2}*Pitslre*[EY22469] | BDSC | 22561 | BDSC_22561 |
| CG5182 | *Pk34A* | *GSK3A* | UAS-*Pk34A* RNAi | VDRC | 101146 | not available |
| CG4379 | *Pka-C1* | *PRKACA* | *Pka-C1*[H2] | BDSC | 4101 | BDSC_4101 |
| CG12066 | *Pka-C2* | *PRKACA* | P{TRiP.JF01448}attP2 (*Pka-C2* RNAi) | BDSC | 31656 | BDSC_31656 |
| CG6117 | *Pka-C3* | *PRKX* | Mi{MIC}*Pka-C3*[MI04599] | BDSC | 37962 | BDSC_37962 |
| CG15862 | *Pka-R2* | *PRKAR2A* | Mi{MIC}*Pka-R2*[MI00092] | BDSC | 30617 | BDSC_30617 |
| CG6622 | *Pkc53E* | *PRKCA* | P{EPgy2}*Pkc53E*[EY14093] | BDSC | 20790 | BDSC_20790 |
| CG1954 | *Pkc98E* | *PRKCE* | PBac{WH}*Pkc98E*[f06221] | BDSC | 18950 | BDSC_18950 |
| CG42349 | *Pkcdelta* | *PRKCD* | PBac{RB}*Pkcdelta*[e04408] | BDSC | 18258 | BDSC_18258 |
| CG7125 | *PKD* | *PRKD1* | Mi{ET1}*PKD*[MB00674] | BDSC | 22868 | BDSC_22868 |
| CG3324 | *Pkg21D* | *PRKG2* | Mi{ET1}*Pkg21D*[MB04805] | BDSC | 24228 | BDSC_24228 |
| CG2049 | *Pkn* | *PKN2* | P{PZ}*Pkn*[06736] | BDSC | 12322 | BDSC_12322 |
| CG5974 | *pll* | *IRAK4* | *pll*[2] | BDSC | 3111 | BDSC_3111 |
| CG11420 | *png* | *NEK11* | *png*[1058] | BDSC | 38437 | BDSC_38437 |
| CG12306 | *polo* | *PLK1* | P{PTT-GC}*polo*[CC01326] | BDSC | 51552 | BDSC_51552 |
| CG7028 | *Prp4k* | *PRPF4B* | P{EPgy2}*Prp4k*[EY11156] | BDSC | 20662 | BDSC_20662 |
| CG10673 | *Tcs5* | *TP53RK* | P{TRiP.GL00310}attP2 (*Tcs5* RNAi) | BDSC | 35391 | BDSC_35391 |
| CG7904 | *put* | *ACVR2B* | P{PZ}*put*[10460] | BDSC | 11745 | BDSC_11745 |
| CG8222 | *Pvr* | *FLT1* | PBac{PB}*Pvr*[c02195] | BDSC | 10856 | BDSC_10856 |
| CG2845 | *Raf* | *BRAF* | *Raf*[7] | BDSC | 7338 | BDSC_7338 |
| CG42667 | *rdgA* | *DGKZ* | *rdgA*[KS60] | BDSC | 31426 | BDSC_31426 |
| CG11111 | *rdgB* | *PITPNM2* | P{EPgy2}*rdgB*[EY20869] | BDSC | 22434 | BDSC_22434 |
| CG14396 | *Ret* | *RET* | Mi{MIC}*Ret*[MI07200] | BDSC | 43099 | BDSC_43099 |
| CG2846 | *Rfk* | *RFK* | P{EP}*Rfk*[G4442] | BDSC | 30081 | BDSC_30081 |
| CG40494 | *RhoGAP1A* | *BCR* | Mi{MIC}*RhoGAP1A*[MI00604] | BDSC | 34396 | BDSC_34396 |
| CG11660 | *RIOK1* | *RIOK1* | P{TRiP.HMC04524}attP40 (*RIOK1* RNAi) | BDSC | 57150 | BDSC_57150 |
| CG11859 | *RIOK2* | *RIOK2* | PBac{WH}*RIOK2*[f03457] | BDSC | 85219 | BDSC_85219 |
| CG12559 | *rl* | *MAPK1* | Mi{MIC}*rl*[MI07033] | BDSC | 43794 | BDSC_43794 |
| CG9774 | *rok* | *ROCK1* | *Rok* [2] | BDSC | 6666 | BDSC_6666 |
| CG4926 | *Ror* | *ROR1* | P{TRiP.HMC05341}attP40 (*Ror* RNAi) | BDSC | 62868 | BDSC_62868 |
| CG7156 | *Rps6kc1* | *RPS6KC1* | Mi{MIC}*Rps6kc1*[MI12022] | BDSC | 56646 | BDSC_56646 |
| CG10539 | *S6k* | *RPS6KB1* | P{PZ}*S6k*[07084] | BDSC | 11713 | BDSC_11713 |
| CG17596 | *S6kII* | *RPS6KA3* | P{EP}*S6kII*[G1845] | BDSC | 32601 | BDSC_32601 |
| CG7001 | *S6KL* | *SGK494* | P{EPgy2}*S6KL*[EY06723] | BDSC | 16752 | BDSC_16752 |
| CG7186 | *SAK* | *PLK4* | PBac{PB}*SAK*[c06612] | BDSC | 17774 | BDSC_17774 |
| CG1891 | *sax* | *ACVR1* | *sax*[5] | BDSC | 8785 | BDSC_8785 |
| CG18085 | *sev* | *ROS1* | sev[14] | BDSC | 10546 | BDSC_10546 |
| CG6114 | *sff* | *BRSK1* | Mi{ET1}*sff*[MB06603] | BDSC | 25517 | BDSC_25517 |
| CG2621 | *sgg* | *GSK3A* | *sgg*[1] | BDSC | 4095 | BDSC_4095 |
| CG18247 | *shark* | *SYK* | *Shark*[1] | BDSC | 5865 | BDSC_5865 |
| CG4290 | *Sik2* | *SIK2* | P{EP}*Sik2*[G366] | BDSC | 33449 | BDSC_33449 |
| CG42856 | *Sik3* | *SIK3* | P{EPgy2}*Sik3*[EY14354] | BDSC | 20921 | BDSC_20921 |
| CG32484 | *Sk2* | *SPHK1* | P{SUPor-P}*Sk2*[KG05894] | BDSC | 14133 | BDSC_14133 |
| CG4527 | *slik* | *SLK* | P{SUPor-P}*Slik*[KG04837] | BDSC | 14435 | BDSC_14435 |

| | | | | | | |
|---|---|---|---|---|---|---|
| CG43756 | *Slob* | *PXK* | PBac{GAL4D,EYFP}*Slob*[PL00361] | BDSC | 19468 | BDSC_19468 |
| CG2272 | *slpr* | *MAP3K9* | Mi{ET1}*slpr*[MB03655] | BDSC | 24455 | BDSC_24455 |
| CG8485 | *Snrk* | *SNRK* | PBac{PB}*Snrk*[c00431] | BDSC | 85941 | BDSC_85941 |
| CG42347 | *sqa* | *MYLK3* | PBac{WH}*sqa*[f01512] | BDSC | 18446 | BDSC_18446 |
| CG44128 | *Src42A* | *FRK* | *Src42A*[myri] | BDSC | 6453 | BDSC_6453 |
| CG7524 | *Src64B* | *FYN, SRC* | PBac{PB}*Src64B*[c04709] | BDSC | 11578 | BDSC_11578 |
| CG8174 | *SRPK* | *SRPK2* | P{EPgy2}*SRPK*[EY03876] | BDSC | 15685 | BDSC_15685 |
| CG11489 | *srpk79D* | *SRPK3* | PBac{PB}*Srpk79D*[c00270] | BDSC | 10036 | BDSC_10036 |
| CG10522 | *sti* | *CIT* | P{SUPor-P}*sti*[KG01697] | BDSC | 13471 | BDSC_13471 |
| CG40293 | *Stlk* | *STRADA* | P{TRiP.JF02614}attP2 (*Stlk* RNAi) | BDSC | 27305 | BDSC_27305 |
| CG44162 | *Strn-Mlck* | *MYLK2* | PBac{PB}*Strn-Mlck*[c02860] | BDSC | 11089 | BDSC_11089 |
| CG17603 | *Taf1* | *TAF1* | P{EP}*Taf1*[EP421] | BDSC | 17281 | BDSC_17281 |
| CG18492 | *Tak1* | *MAP3K7* | *Tak1*[179] | BDSC | 26275 | BDSC_26275 |
| CG31421 | *Takl1* | *MAP3K7* | *Takl1*[c04375] | BDSC | 11414 | BDSC_11414 |
| CG4803 | *Takl2* | *MAP3K7* | P{XP}*Takl2*[d10454] | BDSC | 19330 | BDSC_19330 |
| CG14217 | *Tao* | *TAOK1* | *Tao*[EP1455] | BDSC | 11458 | BDSC_11458 |
| CG6535 | *tefu* | *ATM* | *tefu*[atm-6] | BDSC | 8626 | BDSC_8626 |
| CG7525 | *Tie* | *TIE1* | Mi{MIC}*Tie*[MI02904] | BDSC | 37331 | BDSC_37331 |
| CG14026 | *tkv* | *BMPR1A* | *tkv*[8] | BDSC | 34509 | BDSC_34509 |
| CG34412 | *tlk* | *TLK2* | P{lacW}*Tlk*[G0054] | BDSC | 11593 | BDSC_11593 |
| CG5092 | *Tor* | *MTOR* | *mTor*[DeltaP] | BDSC | 7014 | BDSC_7014 |
| CG1389 | *tor* | *TIE1* | *tor*[1] | BDSC | 1574 | BDSC_1574 |
| CG5408 | *trbl* | *TRIB2* | P{SUPor-P}*trbl*[KG02308] | BDSC | 13203 | BDSC_13203 |
| CG8637 | *trc* | *STK38* | *trc*[1] | BDSC | 1135 | BDSC_1135 |
| CG14305 | *Tssk* | *TSSK1B* | P{EPg}*Tssk*[HP30350] | BDSC | 22102 | BDSC_22102 |
| CG9746 | *Vps15* | *PIK3R4* | Mi{MIC}*Vps15*[MI05805] | BDSC | 42118 | BDSC_42118 |
| CG4488 | *wee* | *WEE1* | *Wee1*[ES1] | BDSC | 5833 | BDSC_5833 |
| CG10776 | *wit* | *BMPR2* | *wit*[A12] | BDSC | 5173 | BDSC_5173 |
| CG8789 | *wnd* | *MAP3K13* | Mi{PT-BM.2}*wnd*[MI00494-BM.2] | BDSC | 39655 | BDSC_39655 |
| CG7177 | *Wnk* | *WNK1* | P{EPgy2}*Wnk*[EY10165] | BDSC | 16970 | BDSC_16970 |
| CG31127 | *Wsck* | *PDGFRL* | P{TRiP.GL00134}attP2 (*Wsck* RNAi) | BDSC | 35247 | BDSC_35247 |
| CG12072 | *wts* | *LATS1* | *wts*[x1] | BDSC | 44251 | BDSC_44251 |
| CG1973 | *yata* | *SCYL1* | P{TRiP.HMS00786}attP2 (*yata* RNAi) | BDSC | 32986 | BDSC_32986 |

**Table S3. Core driver alterations in human bladder cancer cell lines.**
*EGFR* expression and *TP53* and *RB1* alteration status in 5637, UM-UC-3, HT-1376, and HT-1197 cells. Cell-line annotations were compiled from Cellosaurus and cross-referenced with public datasets including the Cancer Dependency Map (DepMap) and Cancer Cell Line Encyclopedia (CCLE).

| Cell line | *EGFR* expression log2(TPM+1) | *TP53* alteration | *TP53* predicted product | *RB1* alteration | *RB1* predicted product |
|---|---|---|---|---|---|
| 5637 | 6.42 | c.839G>C; p.Arg280Thr (AGG→ACG) | Arg to Thr | c.975T>A; p.Tyr325Ter (TAT→TAA) | Tyr to Ter |
| UM-UC-3 | 5.31 | c.338T>G; p.Phe113Cys (TTC→TGC) | Phe to Cys | None detected | No alteration detected |
| HT-1376 | 6.94 | c.749C>T; p.Pro250Leu (CCG→CTG) | Pro to Leu | c.2104C>T; p.Gln702Ter (CAG→TAG) | Gln to Ter |
| HT-1197 | 5.94 | c.1094A>G; p.His365Arg (CAC→CGC) | His to Arg | None detected | No alteration detected |

**Table S4. Culture media for human bladder cancer cell lines.**

RRID, Research Resource Identifier; FBS, fetal bovine serum. Final concentrations of FBS and penicillin/streptomycin are indicated.

| Cell line | RRID | Medium | FBS | Penicillin/streptomycin |
|---|---|---|---|---|
| 5637 | CVCL_0126 | Roswell Park Memorial Institute 1640 | 10% | 1% |
| UM-UC-3 | CVCL_1783 | Eagle's Minimum Essential Medium | 10% | 1% |
| HT-1376 | CVCL_1292 | Eagle's Minimum Essential Medium | 10% | 1% |
| HT-1197 | CVCL_1291 | Eagle's Minimum Essential Medium | 10% | 1% |

**Table S5. Significant suppressors identified in the dominant modifier screen.**

| *Drosophila* gene | Human ortholog |
|---|---|
| *Taf1* | *TAF1* |
| *Tor* | *MTOR* |
| *Asator* | *TTBK2* |
| *gskt* | *GSK3B* |
| *Pka-C3* | *PRKX* |
| *Dyrk3* | *DYRK3* |
| *Tssk* | *TSSK1B* |
| *Pi3K92E* | *PIK3CA* |
| *trbl* | *TRIB2* |
| *Ask1* | *MAP3K15* |
| *Atg1* | *ULK1* |
| *rl* | *MAPK1* |
| *Egfr* | *EGFR* |
| *hppy* | *MAP4K3* |
| *CG31183* | *NPR2* |
| *htl* | *FGFR* |
| *Pkn* | *PKN2* |
| *Dsor1* | *MEK* |
| *sgg* | *GSK3A* |
| *Pitslre* | *CDK11B* |
| *Gyc76C* | *NPR1* |
| *pll* | *IRAK4* |
| *dop* | *MAST1* |
| *dock* | *NCK1* |
| *CG12069* | *PRKACA* |
| *InR* | *INSR* |
| *IKKbeta* | *CHUK* |
| *CaMKII* | *CAMK2D* |
| *Bub1* | *BUB1* |
| *tkv* | *BMPR1A* |
| *CG1344* | *SCYL3* |
| *hep* | *MAP2K7* |
| *ksr* | *KSR* |
| *Raf* | *BRAF* |
| *Ret* | *RET* |
| *tefu* | *ATM* |
| *CG3277* | *STYK1* |
| *gish* | *CSNK1G3* |
| *Rfk* | *RFK* |

| | |
|---|---|
| *CG42637* | *NPR1* |
| *hpo* | *STK3* |
| *ND-42* | *NDUFA10* |
| *rdgB* | *PITPNM2* |
| *CG8173* | *PBK* |
| *cdi* | *TESK2* |
| *Pka-R2* | *PRKAR2A* |
| *GckIII* | *MST4* |
| *Snrk* | *SNRK* |
| *CG1227* | *STK16* |
| *Nak* | *AAK1* |
| *bt* | *TTN* |
| *wts* | *LATS1* |
| *msn* | *MAP4K4* |
| *Prp4k* | *PRPF4B* |
| *JIL-1* | *RPS6KA5* |
| *CG5790* | *CDC7* |
| *hipk* | *HIPK2* |
| *CkIIalpha* | *CSNK2A1* |
| *slik* | *SLK* |
| *CG4629* | *NIM1* |
| *Pkc98E* | *PRKCE* |
| *Cdk2* | *CDK2* |
| *CG8878* | *VRK3* |
| *Drak* | *STK17A, STK17B* |
| *FER* | *FER* |
| *Gprk2* | *GRK5* |
| *Adck5* | *ADCK5* |
| *trc* | *STK38* |
| *fu* | *STK36* |
| *CG3216* | *NPR1* |
| *Ddr* | *DDR2* |
| *Csk* | *CSK* |
| *Takl2* | *MAP3K7* |
